# Regulation of cell dynamics by rapid transport of integrins through the biosynthetic pathway

**DOI:** 10.1101/2022.07.12.498931

**Authors:** Martina Lerche, Mathilde Mathieu, Lene Malerød, Nina Marie Pedersen, Hellyeh Hamidi, Megan Chastney, Bart Marlon Herwig Bruininks, Shreyas Kaptan, Guillaume Jacquemet, Ilpo Vattulainen, Pere Roca-Cusachs, Andreas Brech, Franck Perez, Gaelle Boncompain, Stéphanie Miserey, Johanna Ivaska

## Abstract

Cells sense and respond to the extracellular matrix (ECM) milieu through integrin proteins. Integrin availability on the plasma membrane, regulated by endosomal receptor uptake and recycling, has been extensively studied and regulates cell dynamics in various normal and pathological contexts^1–5^. In contrast, the role of integrin transport through the biosynthetic pathway has been considered primarily as a mechanism to replenish the receptor pool and too slow to influence cell dynamics^6^. Here, we adopted the RUSH (Retention Using Selective Hooks) assay to synchronize integrin anterograde transport from the endoplasmic reticulum (ER), allowing spatial and temporal analysis of newly synthesized receptor traffic. We observe that the delivery of new integrins to the plasma membrane is polarized in response to specific ECM ligands, facilitates integrin recruitment specifically to the membrane-proximal tip of focal adhesions (FA) and contributes to cell protrusion and FA growth. We explain the augmented adhesion growth using a computational molecular clutch model^7^, where increased integrin availability drives recruitment of additional integrins. Notably, a subset of newly synthesized integrins undergo rapid traffic from the ER to the cell surface to facilitate localized cell spreading, seemingly bypassing the Golgi. This unconventional secretion is dependent on cell adhesion and mediated by Golgi reassembling stacking proteins (GRASPs) association with the PDZ-binding motif in the integrin α5 cytoplasmic tail. This spatially targeted delivery of integrins through the biosynthetic pathway may propel cell dynamics by rapidly altering adhesion receptor availability, providing cells with an additional degree of plasticity to respond to their environment.

---

Numerous studies have investigated cell adhesion dynamics as a function of integrin diffusion along the plasma membrane^8^, integrin activation and tethering to the cytoskeleton^8–10^, proteolytic cleavage of adhesion components, microtubule-dependent adhesion turnover and integrin endocytosis from the plasma membrane^11–13^ and recycling of endocytosed integrins back to the cell surface^14–16^. These studies, however, have focused on mature integrin plasma membrane recruitment and traffic. In contrast, the role of integrin transport through the biosynthetic pathway is virtually unexplored and overlooked in regulating cell dynamics, owing to lack of suitable methodology. Biochemical metabolic labelling^6,17^ have classed integrin synthesis and maturation through the biosynthetic pathway as a slow and steady means to replenish the integrin pool.

We employed the RUSH (retention using selective hooks) system^18^, which has been previously used to synchronize and study post-Golgi anterograde trafficking of a variety of cargos^18–20^. We employed the method to integrins to control retention and release from the ER (Extended Data Fig. 1) and to explore, for the first time, the context-dependent traffic of newly synthesized integrins and its implications in cell adhesion and dynamics in real time. Integrin α5β1 is the main fibronectin receptor in many cell types and has been widely studied in the field. Thus far, integrin α5β1 has only been tagged on the C-terminal tail potentially interfering with some established protein-protein interactions^21,22^ To identify a suitable alternative tagging site on the receptor’s ectodomain, we examined the published crystal structure of the integrin α5β1 headpiece^23^. Given that the N-terminus of the α5 polypeptide is localized between the two integrin subunits, away from the fibronectin ligand binding site, we inserted the IL-2 signal peptide, the streptavidin binding peptide (SBP) and enhanced green fluorescent protein (EGFP) at the N-terminus to generate an SBP-EGFP-integrin α5 construct (henceforth referred to as RUSH-α5) (Fig. 1a, Movie 1, Extended Data Fig. 1). Computational modelling revealed that while the flexible EGFP C-terminal region (plus linker) is just long enough to allow direct contact between EGFP and the fibronectin ligand binding site (Extended Data Fig. 2; Movie 2 and 3), EGFP is stably positioned and cannot be displaced unless unphysical forces are applied. Atomistic simulations were consistent with these observations (Movies 4-6; see methods for details). Thus, EGFP tagging of the integrin α5 ectodomain does not interfere with fibronectin binding or α5β1 subunit heterodimerization (Extended Data Fig. 2, and Movies 1-6). In cells, RUSH-α5 was retained in the ER when co-expressed with an ER hook protein composed of streptavidin fused to the ER-retrieval motif KDEL (Fig. 1b) and released upon biotin addition to be transported to the plasma membrane (Fig. 1b; Movie 7; Extended Data Fig. 1 and Extended Data Fig. 3a).

**Figure 1.**
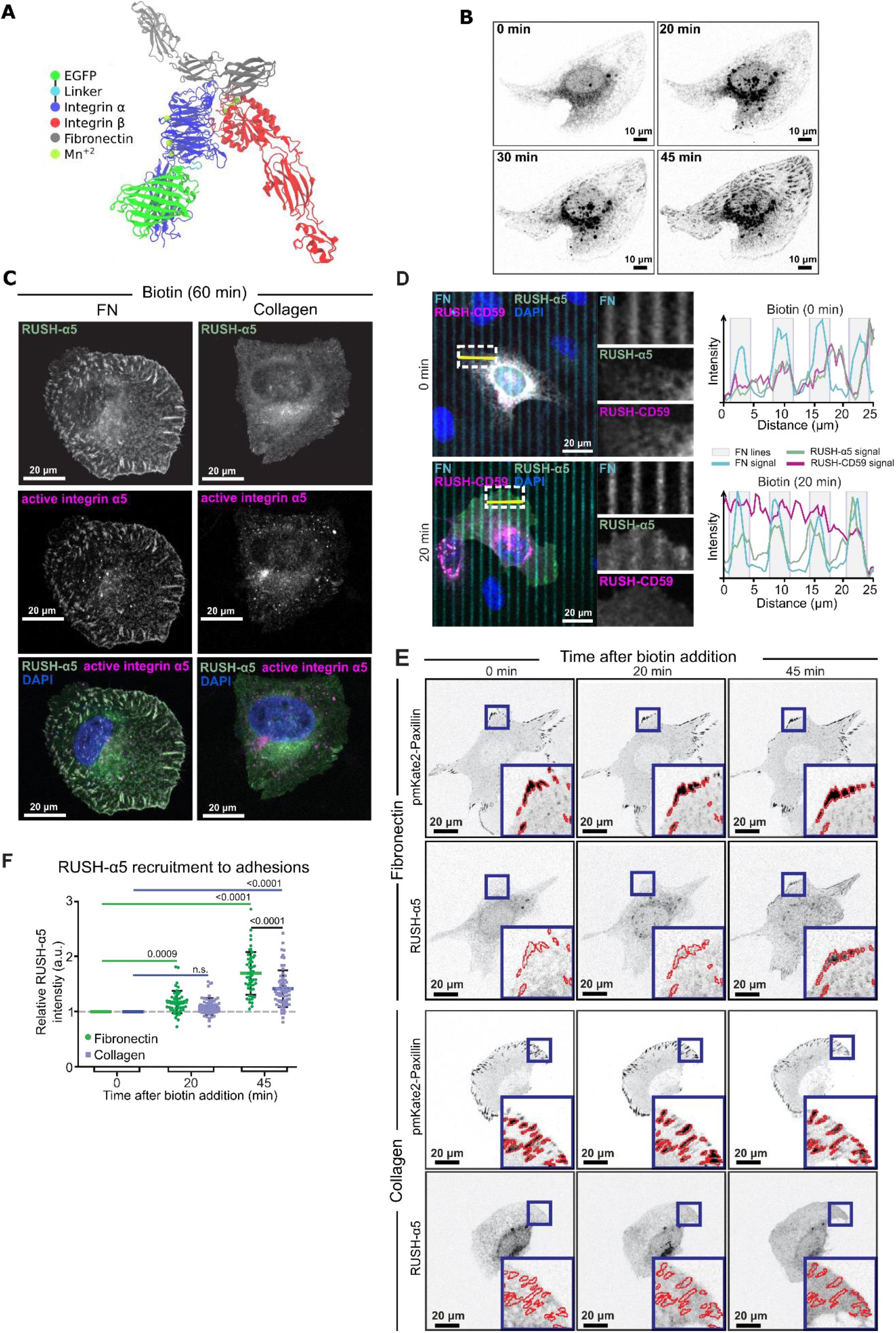
RUSH-α5 delivery to the plasma membrane is spatially regulated by the ECM. **a)** Model of RUSH-α5 (EGFP-integrin α5) - integrin-β1 heterodimer based on (PDB: 7NWL) structure of the heterodimer bound to fibronectin. (see also Movie 1). **b)** Representative immunofluorescence images of RUSH-α5 (SBP-EGFP-integrin α5)-expressing of a time lapse imaged U2OS cell plated on fibronectin ± biotin treatment for the indicated times (see also Movie 7). **c)** Representative immunofluorescence images of RUSH-α5 (green) and active integrin α5β1 (magenta; SNAKA51 antibody) in RUSH-α5-expressing cells plated on fibronectin or collagen + biotin (60 min). Nuclei (blue) were co-labelled. **d)** Representative images of RUSH-α5 and RUSH-CD59 release in cells co-expressing both constructs and plated on dual-coated micropatterns (alternating fibronectin coating (cyan) and collagen-peptide (GFOGER) (non-fluorescent) lines). Nuclei (blue) are co-labeled. Intensity line profiles generated across the yellow line are displayed relative to the position of the fibronectin-coated micropattern lines. White insets represent regions of interest (ROIs) that are magnified for each channel. FN: fibronectin. **e)** Representative immunofluorescence images of cells co-expressing RUSH-α5 and pmKate2-Paxillin plated on fibronectin or collagen ± biotin treatment for the indicated times. Insets represent ROIs that are magnified and show paxillin-segmented adhesions (red outlines). **f)** Quantification of the relative mean intensity of RUSH-α5 in segmented adhesions/cell ± biotin treatment for the indicated times. Data are mean ± SD; n = 64 cells on collagen, 50 cells on fibronectin, pooled from 3 independent experiments; One-way ANOVA, Holm-Sidak’s multiple comparison test.

In cells, integrin β1 subunits are produced in excess and are transported to the plasma membrane only upon heterodimerization with newly synthesized α-integrins^6,24,25^. To investigate the ability of RUSH-α5 to form functional heterodimers with the β1 subunit, we performed GFP-pulldown of RUSH-α5 from cells with and without biotin addition. The EGFP-tagged RUSH-α5 precipitated endogenous β1 integrin. It interacted with the immature integrin β1 (faster migrating lower band; Extended Data Fig. 3b) before release from the ER (0 min biotin), and progressively with the mature integrin β1 after release from the ER following biotin addition (slower migrating upper band; Extended Data Fig. 3b). Importantly, RUSH-α5, when released from the ER (60 min biotin), localized to fibrillar and focal adhesion-like structures on fibronectin positive for active integrin α5 (detected with active conformation specific SNAKA51 antibody) (Fig. 1c). In contrast, RUSH-α5 displayed a clearly reduced localization to adhesion structures along with a more diffuse localization pattern in cells plated on collagen and treated with biotin (Fig. 1c). The ECM ligand did not dramatically influence integrin delivery and maturation as integrin heterodimer maturation (higher ratio of mature to immature integrin β1) was only slightly faster in cells plated on fibronectin compared to cells plated on collagen at 20 and 40 min biotin addition (Extended Data Fig. 3 c-d). Our real-time measurements revealed, however, an interesting and unexpected feature of the kinetics of integrin maturation and delivery. Metabolic labelling studies have indicated receptor maturation kinetics exceeding 1 hour for the total integrin β1 cellular pool^6,17^, which is slower than the α5β1 integrin maturation detected here already 20 minutes of integrin release from the ER (Extended Data Fig. 3c-d). Taken together, these data indicate that the ecto-tagged RUSH-α5 forms a functional heterodimer with endogenous integrin β1, undergoes ligand specific activation on fibronectin and the plasma membrane delivery of newly synthesized integrin is more rapid than previous thought.

To explore the biosynthetic delivery of integrins more closely, we performed time-lapse imaging of the RUSH-α5 release in cells plated on collagen and fibronectin, comparing it with the dynamics of a co-expressed control cargo protein^18^ (glycosylphosphatidylinositol-anchored proteins (GPI-APs) cluster of differentiation 59 (CD59) tagged with SBP and mCherry to use in the RUSH system (henceforth called RUSH-CD59). On both ECMs, RUSH-α5 and CD59 were localized to the ER in the absence of biotin, and following biotin addition were released and transported to the Golgi, residing there for approximately 15 min (Extended Data Fig. 4), in line with previous reports for CD59^18,19^ RUSH-α5 was mostly transported from the Golgi complex after 20 min and recruited to adhesion-like structures on fibronectin whereas on collagen it was diffusely distributed along the plasma membrane (Extended Data Fig. 4 and Movie 8). In contrast, the RUSH-CD59 construct behaved similarly on both fibronectin and collagen (Extended Data Fig. 4 and Movie 8), indicating that the observed differences in RUSH-α5 localization in cells plated on fibronectin and collagen were ligand-receptor specific. This was further validated by plating RUSH-α5 and RUSH-CD59 co-transfected cells on dual-coated micropatterns with alternating lines of fibronectin and GFOGER (a synthetic collagen peptide with high affinity for collagen-binding integrins^26^). Following release (20 min biotin), RUSH-α5 localized predominantly to the cell edges and specifically clustered on the fibronectin lines (Fig. 1d). In contrast, RUSH-CD59 localization was independent of ligand coating. Next, we examined targeting of RUSH-α5 to adhesions by co-expressing pmKate2-paxillin as a focal adhesion marker. RUSH-α5 was recruited to pmKate2-paxillin-positive adhesions already 20 min following release with biotin, and this localization increased over time (Fig. 1e and f). On collagen, RUSH-α5 recruitment to adhesions was significantly lower compared to fibronectin (Fig. 1e and f), with the increase in intensity at the later timepoint most likely reflecting the increased presence of the diffused receptor on the membrane.

Thus, our results show that newly synthesized integrin α5 is trafficked in a ligand-dependent manner and recruited to adhesions. We then investigated if the delivery of newly synthesized integrins is polarized. First, we plated RUSH-α5 transfected cells on 9 μm wide collagen or fibronectin micropatterned lines, shown previously to support front-rear cell polarity of integrins^27^ To detect the localization of RUSH-α5 secretion, we prevented receptor diffusion after plasma membrane delivery by employing the selective protein immobilization (SPI)-method^19^ The ECM proteins are coated alongside anti-GFP antibodies which bind to the luminal GFP moiety in RUSH-α5 when the receptor is delivered to the cell surface. Time-lapse imaging revealed a significant increase in RUSH-α5 intensity over time preferentially at the protruding edge of the cell (region of interest 1; ROI1) on fibronectin (Fig. 2a,b) whereas on collagen lines, the difference in RUSH-α5 localization between the two cell edges was modest and appeared only at later time points (Fig. 2a, c). In line with these data, in unconstrained cells RUSH-α5 delivery was also polarized on fibronectin, occurring significantly more in protruding regions of the cells, whereas on collagen RUSH-α5 intensity increased slowly in retracting and protruding regions (protruding or retracting areas of the cells defined based on spatiotemporal track maps^28^ generated from paxillin images) (Fig. 2d, e). These data indicate that plasma membrane delivery of newly synthesized integrins is sensitive to ECM ligand engagement and cellular front-rear polarity.

**Figure 2.**
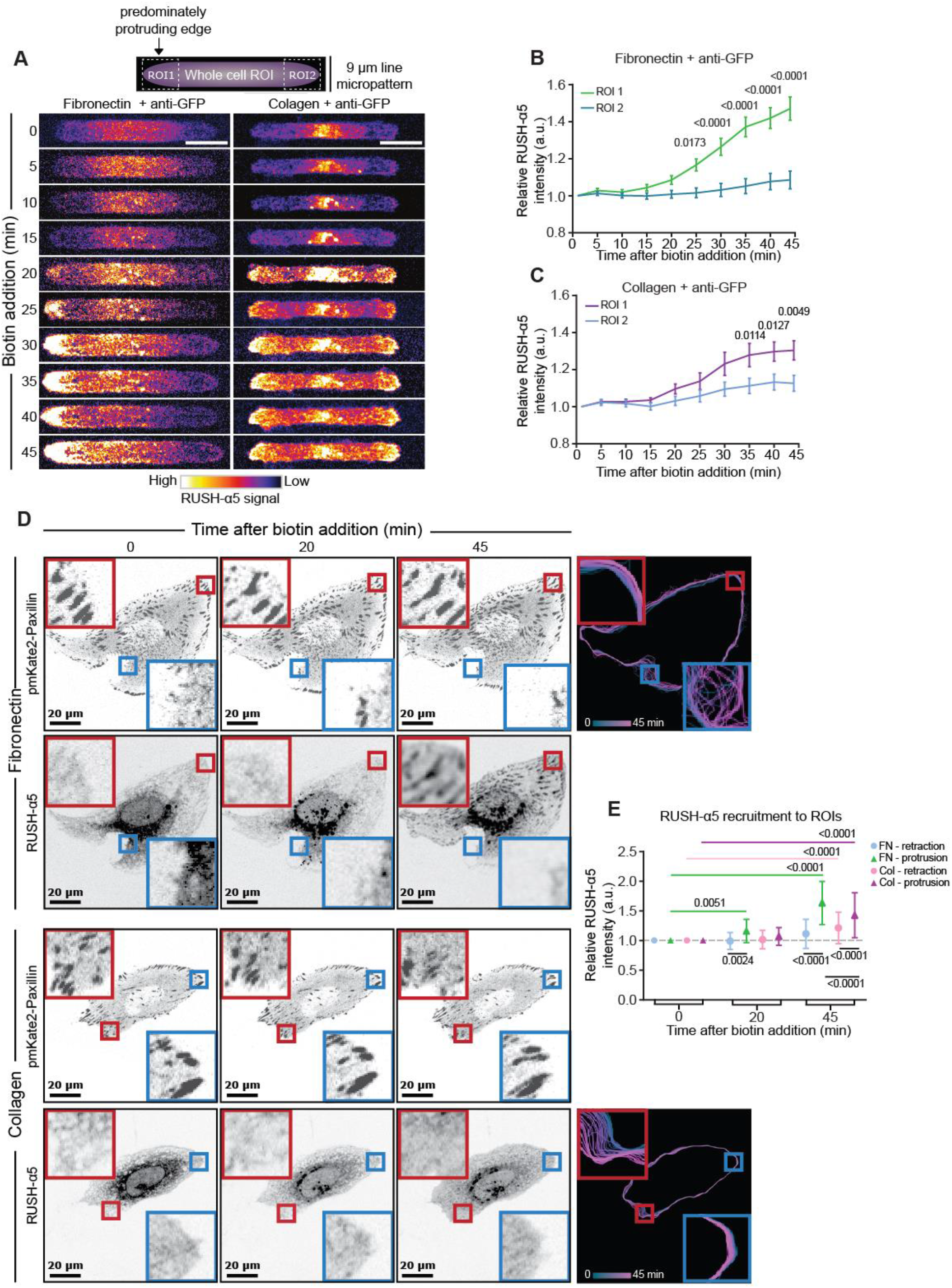
Polarized delivery of newly synthesized integrin to the cell protruding edge. **a-c)** Schematic of the two ROIs chosen for analysis of RUSH-α5 intensity in cells plated on 9 μm wide micropatterns coated with fibronectin and anti-GFP or collagen and anti-GFP and treated with biotin for the indicated times. Representative intensity coded images (a) and quantification of RUSH-α5 release on fibronectin (b) and collagen (c) (normalized first to the total intensity of the cell and then to 0 min biotin) are shown. Data are mean ± SEM. **d)** Representative images and spatiotemporal track maps of cell edge contours over time in cells expressing RUSH-α5 ± biotin treatment for the indicated times. Red insets represent protruding ROIs that are magnified. Blue insets represent retracting ROIs that are magnified. Spatiotemporal track maps: blue colors represent early time points and magenta colors represent late time points in the time-lapse series. **e)** Quantifications of RUSH-α5 intensity in ROIs (retracting or protruding areas determined from spatiotemporal track maps). Data are mean +/− SD. Scale bars: 20 μm. b, c) N = 33 cells on fibronectin and 38 cells on collagen, pooled from three independent experiments, Two-Way ANOVA, Sidak’s multiple comparison test. e) N = 53 cells on collagen, 49 cells on fibronectin, pooled from three independent experiments, One-way ANOVA, Holm-Sidak’s multiple comparisons test.

While RUSH-α5 was predominantly trafficked to the plasma membrane via the Golgi complex conventional secretion pathway, a process that takes more than 20 min, surprisingly some RUSH-α5-positive vesicles were evident earlier (around 10 min) within the vicinity of the plasma membrane (Fig. 3a, Movie 9). This unexpected early plasma membrane delivery was also apparent on the fibronectin line micropatterns (Fig. 3b) and not observed with the RUSH-CD59 construct, which localized in the Golgi complex or the ER (Fig. 3b), indicating that the early secretion was not a general feature for all RUSH cargo. Cell surface delivery of RUSH-α5 was also detected with flow cytometry 15 min after biotin addition with a steady increase up to 1h (Fig. 3c). Live imaging of RUSH-α5 together with an ER marker (ERoxBFP) revealed RUSH-α5 puncta emanating from the ER and being trafficked in close proximity to focal adhesions at very early time points (Movie 9).

**Figure 3.**
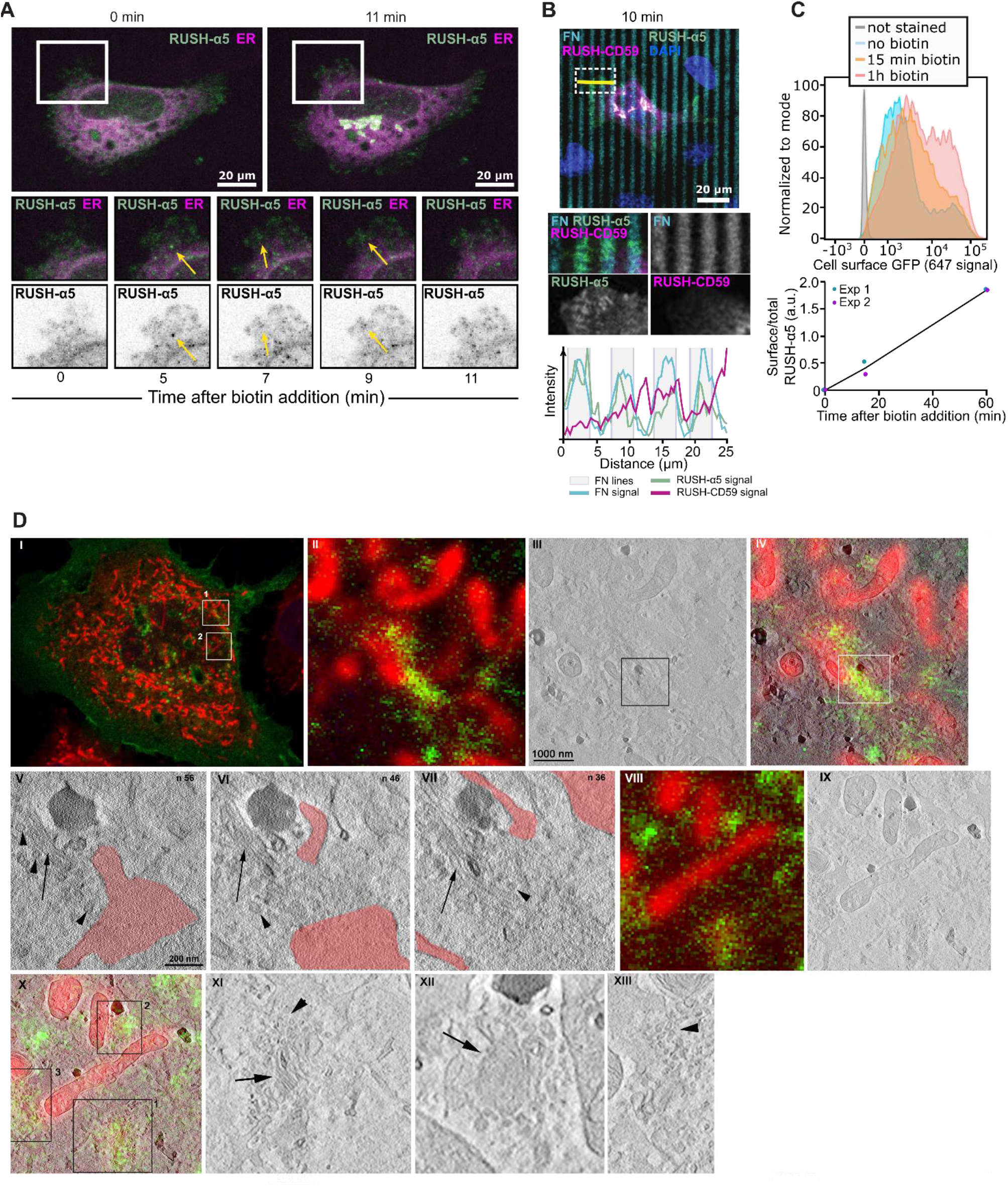
Early delivery of RUSH-α5 from the ER. **a)** Representative immunofluorescence images of cells co-expressing RUSH-α5 (green) and the ER marker ERoxBFP plated on fibronectin (10 μg/ml) ± biotin treatment for the indicated times. Arrows indicate rapidly budding RUSH-α5-positive vesicles adjacent to cell protrusions (≤ 15 minutes after release). (see also Movie 9). **b)** Representative images of RUSH-α5 and RUSH-CD59 release in cells co-expressing both constructs and plated on dual-coated micropatterns (alternating fibronectin coating (cyan) and collagen-peptide (GFOGER) (non-fluorescent) lines). Nuclei (blue) are co-labeled. Intensity line profiles generated across the yellow line are displayed relative to the position of the fibronectin-coated micropattern lines. White insets represent regions of interest (ROIs) that are magnified for each channel. FN: fibronectin. **c)** Flow cytometry analysis of cell-surface RUSH-α5 levels (detected with the anti-GFP-AF647 antibody) in RUSH-α5-expressing U2OS cells ± biotin. Representative histograms and quantification of cell surface GFP (ratio of the geometric means of the surface signal divided by the total GFP signal, normalized by subtracting the 0 min value) are shown. **d)** Ultrastructural analysis of integrin exit sites after 10 min of biotin addition using the RUSH assay. Cells were fixed and confocal images were obtained using a Zeiss LSM780 microscope. Structures positive for RUSH-α5 (green) were identified and studied by correlative light/electron microscopy using STEM tomography with a Thermo Scientific^TM^ Talos^TM^ F200C. MitoTracker Red stained mitochondria (red) were used as landmarks for correlation. Single optical section of a transfected cell is shown in (I) (optical section no. 2, 250 nm thickness) with regions of interest (ROI) marked 1 and 2. ROI1 is further magnified in panel II, with the corresponding area imaged by STEM-tomography in III and both overlaid in IV. Several tomogram slices of ROI III and IV are presented in V, VI and VII, representing an approximate distance of 31 nm between them. ROI2 from I shows additional examples of RUSH-α5 positive areas which are magnified in VIII with a corresponding slice from STEM-tomography in IX and the overlay in X. These areas often consist of small tubular elements (arrows in XI (ROI1 in X) and XII (ROI2 in X) and vesicular clusters were frequently observed (arrowheads in XI, XII and XIII (ROI3 in X)) in close proximity between ER-sheets and tubules. Scale bars as indicated.

Correlative light/electron microscopy (CLEM) experiments using the RUSH-α5, released with biotin for 10 minutes, allowed us to further characterize the ultrastructure features of the compartment involved in early secretion of integrin from the ER. There was marked accumulation of GFP signal in a tubular-vesicular compartment in close proximity to the plasma membrane, which often seem to be tightly associated with ER-sheets (Fig. 3d). However due to limits of resolution in our STEM-tomography approach we could not resolve whether there are continuities between the ER and the integrin-positive structures, or whether there are possible budding profiles associated with the ER.

The observed rapid delivery of integrins to the plasma membrane prompted us to hypothesize that a small proportion of integrins could undergo unconventional protein secretion (UPS), a process where secretory proteins are transported from the ER to the plasma membrane without entering the Golgi complex^29^. Conventional Golgi secretion is inhibited by Golgicide A^30^. Therefore, we next treated cells with Golgicide A to investigate the relative contribution of the Golgi complex to RUSH-α5 traffic. As expected, the delivery of the majority of RUSH-α5, 25 min post-release, was significantly inhibited by Golgicide A (Fig. 4a, b). However, both control and Golgicide A-treated cells showed a small initial increase in RUSH-α5 recruitment to protruding areas of the cell and to adhesions 15 min after release (Fig. Fig. 4a, b). These data indicate that a small fraction of newly synthesized integrins are secreted to focal adhesions via a mechanism that bypasses conventional Golgi secretion.

**Figure 4.**
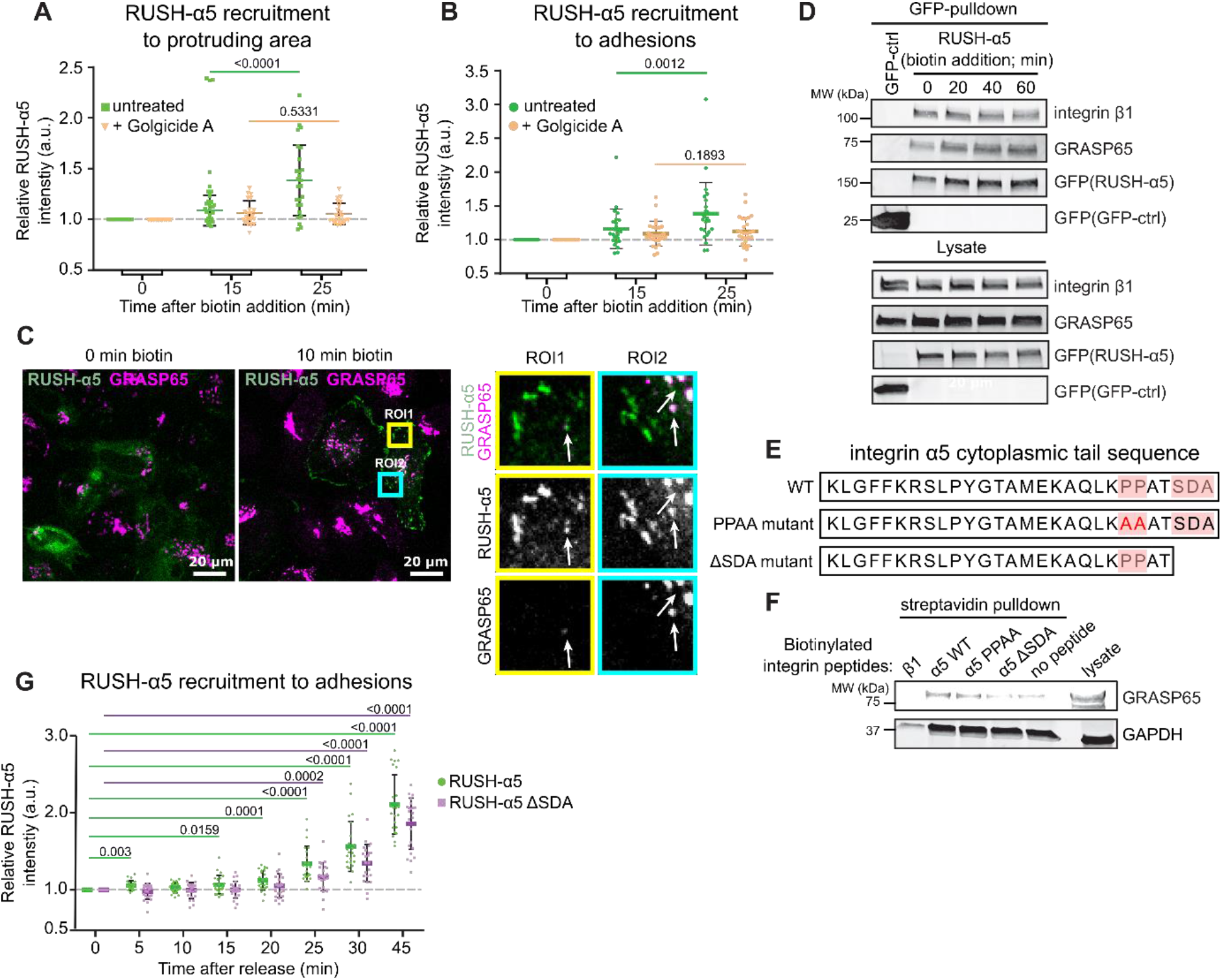
Golgi bypass early delivery of RUSH-α5 is PDZ-motif dependent. **a,b)** Quantification of relative RUSH-α5 recruitment to protruding areas (a) or adhesions (b) in cells expressing RUSH-α5 ± biotin treatment for the indicated times with or without Golgicide A (10 μM). Data are mean ± SD. **c)** Representative image of cells expressing RUSH-α5 (green) and stained for endogenous GRASP65 (magenta) plated on fibronectin ± biotin treatment for the indicated times. **d)** Representative immunoblot of GFP-pulldowns from RUSH-α5 or GFP control transfected cells plated on fibronectin and probed for GFP and endogenous GRASP65. N= 3 independent experiments. **e)** Amino acid sequence of the integrin α5 tail highlighting the canonical PDZ-binding motif (SDA) and the two proline residues critical for the formation of the non-canonical PDZ-binding motif. The mutations of these sites used in our experiments are indicated below. **f)** Representative streptavidin pulldowns of the indicated biotinylated recombinant integrin peptides incubated with cell lysates collected from CHO cells overexpressing GRASP65-GFP. Representative immunoblots probed for GRASP65 and GAPDH are shown. N=3 independent experiments. **g)** Quantification of RUSH-α5 or RUSH-α5 ΔSDA recruitment in adhesions ± biotin treatment for the indicated times. All data are mean +/− SD. Scale bars: 20 μm. a,b) One sample t-test. a) N=26 cells RUSH-α5, N=22 cell RUSH-α5 Golgicide A, pooled from 3 independent experiments. b) N=24 cells RUSH-α5, N=27 cell RUSH-α5 Golgicide, pooled from 3 independent experiments. N=2 independent experiments. g) One sample t-test. N=23 cells RUSH-α5, N= 23 cells RUSH-α5 ΔSDA, pooled from two independent experiments.

Thus far, Golgi bypass traffic of integrins has been reported only for the integrin αPS1 subunit upon mechanical stress at specific stages of Drosophila follicle epithelium development^31^. This prompted us to investigate whether early secretion of newly synthetized integrins in mammalian cells is linked to cell adhesion and adhesion-induced mechanics. First, we explored whether pre-existing endogenous integrin α5 adhesions are involved. We knocked-out endogenous integrin α5 (ITGA5) (Extended Data Fig. 5a-d) and compared RUSH-α5 release in ITGA5 wild-type (WT) and knockout (KO) cells at 15 and 60 min biotin addition (please note that U2OS cells have other fibronectin-binding integrins in addition to integrin α5 and thus the KO cells adhere to fibronectin similarly to control cells). RUSH-α5 delivery to the cell surface was the same between the ITGA5 WT and KO clones (Extended Data Fig. 6a). Thus, the early secretion of newly synthetized integrin α5 is not dependent on the localization of the endogenous protein already at the cell surface. However, the early delivery of RUSH-α5 to the plasma membrane was dependent on cell-ECM adhesion as we did not detect RUSH-α5 at the cell surface after 15 min biotin addition in suspension cells (Extended Data Fig. 6b). Adhesion was not required for slower integrin secretion (60 min biotin) via the conventional secretion pathway (Extended Data Fig. 6b). These data are consistent with cell adhesion and perhaps active spreading/protrusions on rigid surfaces acting as necessary triggers for early secretion of integrin α5 to the cell surface. Even though the early delivery of RUSH-α5 as such was not dependent on endogenous integrin α5, polarized RUSH-α5 delivery to cell protruding areas was significantly higher in WT compared to ITGA5 KO cells (Extended Data Fig. 6c), suggesting that polarized delivery of the newly synthesized integrin is orchestrated by existing adhesions and contributes to rapid alterations in cell shape.

The two mammalian homologues GRASP55 and GRASP65 mediate UPS of transmembrane proteins via PDZ-domain-mediated interactions with cargo proteins^32,33^. We stained endogenous GRASP65 in cells at various time points after RUSH-α5 release and observed small RUSH-α5- and GRASP-positive puncta in close proximity to the cell periphery 10 min post release (Fig. 4c). Integrin α5 cytoplasmic tail harbors two distinct PDZ-binding motifs in its cytoplasmic domain: a classical C-terminal Ser-Asp-Ala (SDA) sequence^34^ and a non-canonical motif generated by two prolines (PP) that induce an internal β hairpin structure to function as a PDZ recognition motif^21^. To test whether integrin α5 associates with GRASP we performed pulldown experiments. RUSH-α5 pulldown with anti-GFP nanobody beads coprecipitated endogenous GRASP65 in cells (Fig. 4d). Pull-down experiments with biotinylated peptides corresponding to the C-terminal part of the integrin α5 WT tail or integrin α5 tails with mutations in the non-canonical PDZ-binding motif (PPAA peptide) or deletion of the canonical PDZ-binding motif (ΔSDA) (Fig. 4e,f) indicated that GRASP65 associates with integrin α5 via the SDA sequence (Fig. 4f), in accordance with the ability of GRASP65 to associate with ER resident cargo to facilitate UPS and regulate N-linked glycosylation in the ER^35^.

We then explored the involvement of GRAPS and the integrin tail in the early delivery of RUSH-α5. siRNA-mediated silencing of GRASP65 and GRASP55 (Extended Data Fig. 7a) inhibited RUSH-α5 delivery to fibronectin spots on dual-coated (FN and GFOGER) micropatterns at 10-15 min post release (Extended Data Fig. 7b-c). However, the recruitment of RUSH-α5 to fibronectin dots remained low 25 min after release, possibly due to GRASP silencing interfering with Golgi function^36^. Furthermore, GRASP depletion has been shown to downregulate integrin α5β1 protein levels and could also affect the lifetime of our exogenous construct^37^ To overcome these complications, we generated a RUSH-α5 construct lacking the GRASP65 binding SDA sequence (RUSH-α5-ΔSDA) and performed time-lapse imaging. RUSH-α5-ΔSDA recruitment to adhesions was considerably delayed compared to RUSH WT (25 min versus 5 min) (Fig. 4g). After 45 minutes, RUSH-α5-ΔSDA levels in adhesions remained lower than WT, consistent with unconventional secretion accounting for a small part of the overall integrin biosynthetic delivery in cells.

The endo/exocytic traffic of cell surface integrins controls FA dynamics, size and distribution in cells^13,38–40^, however the role of integrin secretion remains to be explored. CD59, along with several other cargo proteins, undergo anterograde post-Golgi traffic to secretion hotspots adjacent to but discrete from the FAs^19^. We employed dual color TIRF imaging of RUSH-α5 and pmKate2-Paxillin in cells plated on fibronectin to determine whether this is the case for integrins as well. SPI revealed that RUSH-α5 is recruited directly to the FA rather than adjacent hot spots. It was initially delivered to the most membrane proximal area (Area 1) of FA after which it gradually accumulated along the growing adhesion (Fig. 5a-c), indicating RUSH-α5 is specifically delivered to the membrane proximal tips of adhesions.

**Figure 5.**
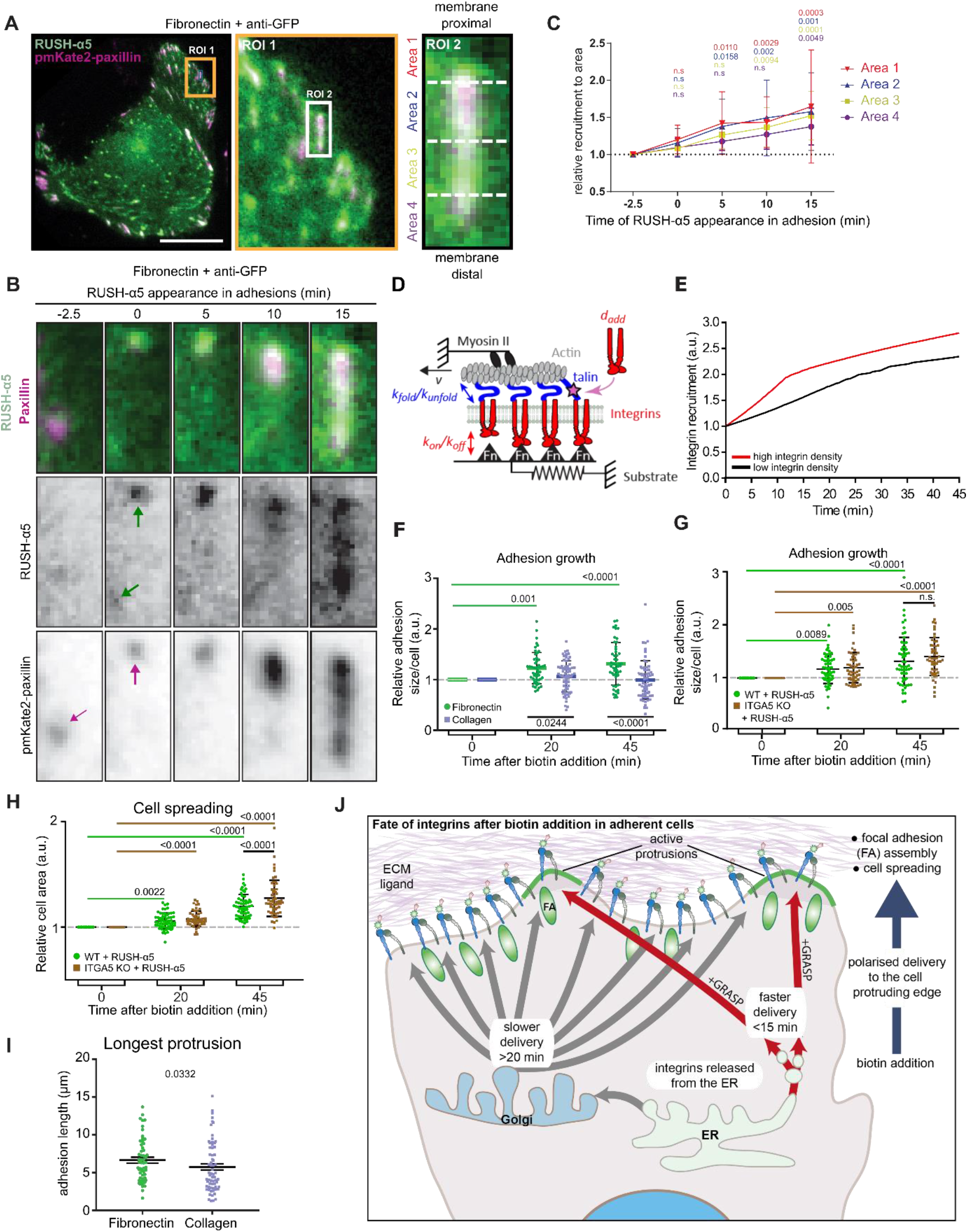
RUSH-α5 is delivered to the tip of adhesions and mediates adhesion growth. **a-b)** Representative image of cells expressing RUSH-α5 and pmKate2-Paxillin plated on fibronectin (10 μg/ml) and anti-GFP (2.5 μg/ml; to trap cell surface released RUSH-α5 at the point of delivery), insets represent ROIs that are magnified. ROI2 represents a focal adhesion demarcated into four equal areas for analysis and is further magnified in b. Scale bar 20 μm. **c)** Quantification of RUSH-α5 intensity in the four areas relative to signal intensity in the respective area 2.5 min prior to RUSH-α5 appearance in adhesions (determined from the time-lapse images). Adhesions close to the cell edge and with a minimum lifetime of 15 min were analyzed and changes of RUSH-α5 intensity were blotted over time in the indicated areas ranging from proximal to distal to the cell edge. **d)** Cartoon showing clutch model elements. Myosin motors pull on actin filaments with a speed *v*. This applies force to a substrate via integrins and adaptor proteins (talin). The effect of force regulates the unbinding rates from integrins to the substrate *(k_off_)* and the folding/unfolding rates of talin (*k_fold_/k_unfold_*). When talin unfolds, adhesion reinforcement is assumed to happen, which is modelled by an increase in integrin density with value *d_add_*. Changes in integrin availability are modelled by changing the parameter *d_add_*. **e)** Model prediction of adhesion growth with time for conditions in which integrin availability is low (*d_add_* = 0.005 integrins/μm^2^) or high (*d_add_* = 0.01 integrins/μm^2^). Adhesion growth (y-axis) is modelled through integrin density, which is plotted normalized to the starting value. **f)** Quantification of adhesion growth in cells expressing RUSH-α5 and plated on fibronectin or collagen ± biotin treatment for the indicated times. Shown are the relative sums of segmented adhesion area/cell at indicated time points. Data are mean ± SD. **g)** Quantification of adhesion growth in WT and ITGA5 KO cells expressing RUSH-α5 ± biotin treatment for the indicated times. Shown are the relative sums of segmented adhesion area/cell at indicated time points. Data are mean ± SD. **h)** Quantification of cell spreading in WT and ITGA5 KO cells expressing RUSH-α5 ± biotin treatment for the indicated times. Data are Mean ± SEM. **i)** Quantifications of the length of the longest protrusion (extending furthest from the initial plasma membrane localization during imaging) formed per cell after 45 minutes of biotin. Data are mean ± SEM. **j)** Schematic depiction of the regulation of cell dynamics by transport of integrins through the biosynthetic pathway. Adhesion and cell spreading dependent delivery of integrin from the ER is detected rapidly after release in cell protrusions. Canonical Golgi-dependent delivery is also polarized to cell protruding areas in an ECM-specific manner and contributes to focal adhesion growth and cell protrusion. (c) One independent experiment 9 adhesions from 6 cells on 2 coverslips. One-way ANOVA, Dunn’s multiple comparison test. (f, g and h) One-way ANOVA, Holm-Sidak’s multiple comparison test. (f) N = 64 cells on Coll, 50 cells on FN, pooled from 3 independent experiments. (g) 57 WT cells and 52 ITGA5 KO cells, (h) 59 WT cells and 55 ITGA5 KO cells, pooled from 3 independent experiments. (i) Mann-Whitney test, N=55 cells on FN, 66 cells on Coll, pooled from 3 independent experiments. ER retention and release of integrin α5 using RUSH (Retention Using Selective Hooks)

To consider the effects of increased integrin delivery on adhesions, we employed a molecular clutch model previously developed to simulate mechanosensitive growth of adhesions^7^. In this model, talin-mediated mechanosensing (through force-induced unfolding) leads to adhesion growth, modelled as an increase in integrin density (Fig. 5d). To understand the effect of integrin delivery, we reasoned that an increased delivery would result in a higher availability of integrins to be incorporated into adhesions. Thus, we modelled integrin delivery by tuning the parameter that sets the increase in integrin density that occurs upon talin unfolding (*d_add_*). Running the model with a base set of parameters taken from previous work (Extended Data Table 1), modifying only the *d_add_* parameter, and running the simulation as a function of time, indicated that delivery of new receptor is predicted to increase adhesion growth (Fig. 5e). In concordance with the model, release of RUSH-α5 significantly increased adhesion area on fibronectin, and had no significant effect on adhesion area on collagen (Fig. 5f). Adhesion growth supported by RUSH-α5 release was also apparent in the ITGA5 KO cells (Fig. 5g), indicating that an increased number of clutches translates to increased adhesion size also when a new type of integrin heterodimer is introduced to the cell surface. The increase in adhesions correlated with increased cell spreading in WT and KO cells (Fig. 5h) and contributed to increased cell dynamics with longer cell protrusions extended in RUSH-α5 released cells plated on fibronectin compared to collagen (Fig. 5i, Extended Data Fig. 8). Taken together, these data indicate that newly synthesized integrin α5 is rapidly localized to the membrane proximal end of focal adhesions, contributing to adhesion growth towards the membrane distal end and facilitating cell protrusion in a spatially defined and ligand-dependent manner.

Altogether, our findings demonstrate that cell adhesion and polarized cell dynamics can be steered by targeted delivery of newly synthesized integrin to the plasma membrane, and that this can occur rapidly, in a localized manner, bypassing the Golgi complex (Fig. 5j). While proteins can be N-glycosylated in the ER^35^, it is plausible that the early secreted pool of integrins is not fully glycosylated but rather in a high-mannose state. N-glycan chains support integrin headpiece opening (activation) and increase integrin ligand-binding affinity^41^. However, high-mannose integrins are also functional in ECM engagement^41^, consistent with early delivery of RUSH-α5 supporting cell protrusions and adhesion growth. The field of cell-matrix adhesion is well-established and there is a wide consensus that endocytosis and recycling of integrins from and to the plasma membrane are essential regulators of cell dynamics, migration and invasion^1–5^. We find here that delivery of fresh integrin, along the biosynthetic pathway, is also operating to determine cell dynamics. Thus, mechanisms regulating integrin secretion are likely to be intertwined with established integrin trafficking pathways in previously unappreciated ways and most likely these mechanisms operate alternately or even simultaneously endowing cells with greater plasticity to adapt to dynamic alterations in their extracellular environments.

## Methods

### Cell culture

CHO cells were grown in Ham’s F12 supplemented with 10 % fetal calf serum (FCS) and 1 % L-glutamine. HEK293-FT and U2OS cells were grown in Dulbecco’s Modified Eagle’s medium (DMEM) supplemented with 10 % fetal calf serum (FCS), 1 % L-glutamine and 1% penicillin/streptomycin. Cells were routinely tested for mycoplasma contamination.

### Plasmids

ERoxBFP was purchased from Addgene (68126) and pmKate2-paxillin from Evrogen (FP323). pEGFP-GRASP65 was a kind gift from Kasper Mygind at Institute Curie. Streptavidin-KDEL_SBP-mCherry-CD59 was generated as previously described^19^ Streptavidin-KDEL_SBP-EGFP-ITGA5 was generated by PCR amplification of human ITGA5 without its signal peptide using Integrin-α5-EGFP template from Patrick Caswell and the following PCR primers: forward 5’-AATTggccggccgTTCAACTTAGACGCGGAGGC-3’ and reverse 5’-AACCttaattaatcaGGCATCAGAGGTGGCTGG-3’. The PCR fragment was then subcloned in the RUSH plasmid Streptavidin-KDEL_ss-SBP-EGFP^18^ using FseI and PacI restriction enzymes. The hook (streptavidin-KDEL) allows anchoring of the SBP-tagged reporter (integrin α5 and CD59) in the ER in the absence of biotin thanks to streptavidin–SBP interaction.

### Transfection

Plasmids of interest were transfected using Lipofectamine 3000, Lipofectamine 2000 (Thermo Fisher Scientific) or jetPRIME^®^ (Polyplus transfection) according to the manufacturer’s instructions. Protein downregulation was carried out with Lipofectamine siRNA Max or Lipofectamine 3000 (ThermoFisher Scientific) according to manufacturer’s instructions. The siRNA used as control (siCTRL) was Allstars negative control siRNA (Qiagen, Cat. No. 1027281). GRASP65 and GRASP55 were downregulated with Flexitube siRNAs (GS64689 and GS26003 respectively, Qiagen) or custom ordered siRNA oligonucleotides targeting GRASP65 and GRASP55 (GRASP65 target sequence: AAG-GCA-CUA-CUG-AAA-GCC-AAU and GRASP55 target sequence: AAC-UGU-CGA-GAA-GUG-AUU-AUU, Qiagen).

### RUSH-α5 transfection and release

Cells grown in 25% confluence were used for transfection. For a 6 cm dish 1×10^5^ cells were reversely transfected with 10 μg RUSH-α5 using Lipofectamine 3000 according to the manufacturers’ protocol. The cells were from this point grown in medium containing 1-2.5 μg/ml streptavidin (S4762, Sigma-Aldrich) to block biotin in the media and transfection reagents. The next day after transfection, cells were used for experiments, either directly by releasing the RUSH in or splitting and seeding the cells beforehand on appropriate surfaces for imaging experiments as described below. The release of the RUSH cargos from the ER-hook was induced by removing the streptavidin supplemented media and addition of biotin supplemented media (3 mM of D-biotin, B4501 Sigma-Aldrich) for the indicated times.

### Immunoprecipitations and immunoblotting

HEK293-FT or U2OS cells expressing GFP-tagged proteins (three 10 cm dish per condition or one 10 cm dish for GFP-controll construct, due to differences in expression efficiency) were washed with cold PBS, harvested in PBS and pelleted. Commercial hydrogels Petrisoft™ 100, Easy Coat™ (PS100-EC-50 and PS100-EC-0.5, Matrigen) were used for samples collected from hydrogels. The cell pellet was resuspended in 200 μl of IP-lysis buffer (40 mM Hepes-NaOH, 75 mM NaCl, 2 mM EDTA, 1% NP40, protease and phosphatase inhibitors) and incubated at +4 °C for 30 min, followed by centrifugation (10,000x g for 10 min, +4 °C). 20 μl of the supernatant was kept aside as the lysate control. The remainder of the supernatant was incubated with GFP-Trap beads (ChromoTek; gtak-20), for 55 min at 4 °C. Finally, immunoprecipitated complexes were washed three times with wash-buffer (20 mM Tris-HCl pH 7.5, 150 mM NaCl, 1 % NP-40) and denatured for 10 min at 95°C in reducing Laemmli buffer before SDS-PAGE analysis under denaturing conditions (4–20 % Mini-PROTEAN TGX Gels). The proteins were then transferred to nitrocellulose membranes (Bio-Rad Laboratories) before blocking with blocking buffer (Thermo, StartingBlock (PBS) blocking, #37538) and PBS (1:1 ratio). The membranes were incubated with primary antibodies diluted in blocking buffer overnight at 4°C. Following this step, membranes were washed three times with TBST and incubated with fluorophore-conjugated secondary antibodies (LI-COR) diluted (1:10,000) in blocking buffer at room temperature for 1 hour. Membranes were scanned using BioRad ChemiDoc MP Gel Analyzer, an infrared imaging system (Odyssey; LI-COR Biosciences) or Azure Sapphire RGBNIR Biomolecular Imager. Primary antibodies used: Mouse anti-CD29 (integrin β1) (610468, BD Biosciences), rabbit anti-GRASP55 (HPA035274, Sigma), rabbit anti-GRASP65 (HPA056283), rabbit anti-GFP (ab290, abcam) and mouse anti-GAPDH (5G4MaB6C5, Bioz). Secondary antibodies used: IRDye^®^ 800CW Donkey anti-Mouse IgG, IRDye^®^ 800CW Donkey anti-Rabbit IgG, IRDye^®^ 680LT Donkey anti-Mouse IgG and IRDye^®^ 680LT Donkey anti-Rabbit IgG, diluted 1:10000 in odyssey blocking buffer (LI-COR).

Pulldown with N-terminally biotinylated peptides (GenScript) was carried out as follows: biotinylated peptides were incubated with streptavidin conjugated Dynabeads (65001, ThermoFisher) for 30 min RT followed by 2 h incubation with supernatant, (prepared in the same way as described above for GFP-immunoprecipitated samples) from EGFP-GRASP65 overexpressing CHO cells, at +4°C. Immunoprecipitated complexes were washed three times with wash-buffer (50 mM Tris-HCl pH 7.5m, 150 mM NaCl and 1 % NP-40) and denatured for 10 min at 95°C in reducing Laemmli buffer before SDS-PAGE analysis.

### Flow Cytometry

Cells were detached on ice, after biotin release in the case of RUSH-ITGA5 transfected cells, with enzyme-free cell dissociation buffer (Gibco 13150016). Pelleted cells were incubated with anti-GFP-AF647 antibody (1:150, Alexa Fluor 647 Mouse Anti-GFP Clone 1A12-6-18, BD Biosciences 565197) or rabbit anti-ITGA5 (1:100, clone EPR7854, Abcam ab150361), in Tyrodes Buffer (10 mM HEPES-NaOH at pH7.5, 137 mM NaCl, 2.68 mM KCl, 1.7 mM MgCl2, 11.9 mM NaHCO3, 5 mM glucose, 0.1% BSA) for 40 min on ice. Cells were washed twice with Tyrodes Buffer. In the case of anti-ITGA5 staining, cells were then incubated with a donkey anti-Rabbit-AF647 (1:400, Invitrogen A31573) for 30 min on ice and washed twice. Cells were fixed for 10 min with 2 % PFA, resuspended in PBS and analyzed with LSRFortessa (BD Biosciences). Data analysis was performed with FlowJo software version 5. For quantifications of the anti-GFP surface stainings, the geometric mean of the anti-GFP antibody signal was divided by the total GFP signal, and the value of this ratio at T0 was subtracted to the ratio at all time points.

### Live cell imaging

Cells were plated and allowed to spread for 2-4 h before imaging on fibronectin or collagen (10 μg/ml) coated coverslips, additional 2.5 μg/ml Alpaca anti-GFP VHH nanobody (gt-250, Chromotec) coating was used in indicated experiments. Time-lapse imaging was performed at 37°C using Spinning-disk confocal 3i (Intelligent Imaging Innovations, 3i Inc) Marianas Spinning disk confocal microscope with a Yokogawa CSU-W1 scanner and back illuminated 10 MHz EMCDD camera (Photometrics Evolve) using 63x/1.4 oil objective. TIRF imaging was carried out using DeltaVision OMX with 60x/1.49 Olympus APO N TIRF objective. Conventional protein secretion was blocked (in indicated experiments) by incubating the cells with 10 μM Golgicide A (G0923, Sigma-Aldrich) 30 minutes prior to imaging. The release of the RUSH cargos was induced by removing the streptavidin supplemented media (1μg/ml streptavidin, S4762, Sigma-Aldrich) and addition of biotin supplemented media (3mM of D-biotin, B4501 Sigma-Aldrich), by using a magnetic imaging chamber with L-shape tubing (CM-B25-1PB, Live Cell Instrument CO., LTD) during the live cell imaging experiments.

### Image analyses

As the intensity of RUSH-α5 varied from cell to cell based on transfection efficiency, relative RUSH-α5 recruitment was measured by normalizing the intensity at indicated time point to the same intensity measurement before release in the same cell. Due to the low exposure time used for image acquiring of pmKate2-Paxillin (to reduce photo toxicity), denoising of paxillin adhesions was carried out using the deep learning CARE2D network^42^ before segmenting the paxillin adhesions when needed. Images of paxillin adhesions were made binary and adhesions larger than 0.6 μm^2^ were segmented and analyzed with the Analyze Particles tool in Image J. The segmented adhesions were saved as ROIs in the ROI manager and used to measure the intensity of RUSH-α5 within the paxillin adhesions. Spatio-temporal track maps of cells were generated based on the RUSH-α5 signal using the QuimP plugin^28^ in ImageJ. Localization of RUSH-α5 to adhesions was studied by drawing a ROI around the whole adhesion area based on the paxillin signal and then dividing the ROI into 4 areas of equal size. RUSH-α5 intensity in the four areas relative to signal intensity in the respective area 2.5 min prior to RUSH-α5 appearance in adhesions (determined from the time-lapse images) was measured. Adhesions close to the cell edge and with a minimum lifetime of 15 min were analyzed and changes of RUSH-α5 intensity were blotted over time in the indicated areas ranging from proximal to distal to the cell edge.

### Immunofluorescence and image acquiring of fixed samples

Cells were plated on ibidi 35 mm μ-dishes coated with 10 μg/ml collagen I or fibronectin. Samples were fixed for 10 min with 4 % PFA followed by permeabilization for 10 min with 0.1 % Triton X-100 in PBS. To block unspecific binding of antibodies cells were incubated in 10 % horse serum (HRS) for 1 h or in 1 M Glycine for 20 min in RT. Primary and secondary antibodies were diluted in 10 % HRS and incubated for 1h in RT. Primary antibodies used: mouse anti-integrin α5 (clone SNAKA51) (MABT201, Millipore), rabbit anti-GRASP55 (HPA035274, Sigma) and rabbit anti-GRASP65 (HPA056283). Secondary antibodies used: Alexa Fluor 568 anti-rabbit (A-11011 Thermo Fisher 1:400), Alexa Fluor 555 anti-mouse (A32727, Thermo Fisher 1:400). F-actin was stained with Phalloidin-Atto 647N (65906, Sigma, 1:400), incubated together with secondary antibodies and nuclei with DAPI (D1306, Life Technologies 1:3000) for 10 min RT after secondary antibody incubation. Samples were imaged using either A) 3i (Intelligent Imaging Innovations, 3i Inc) Marianas Spinning disk confocal microscope with a Yokogawa CSU-W1 scanner and Hamamatsu sCMOS Orca Flash 4.0 camera (Hamamatsu Photonics K.K.) or back illuminated 10 MHz EMCDD camera (Photometrics Evolve) using 63x/1.4 oil objective or 40x/1.1 water objective, or B) Zeiss LSM780 laser scanning confocal microscope using 40x/1.2 water Zeiss C-Apochromat objective.

### Micropatterns

Micropatterns were produced on glass coverslips as described in^43^. Cells were seeded in media with fibronectin deprived serum and allowed to spread on micropatterns for a minimum 2 h. For experiments with double coated micropatterns PLL(20)-g[3.5]-PEG(2)/PEG(3.4)-biotin(50%) (SuSoS) was used to allow for coating of also the non-micropatterned areas: the micropatterned areas were first coated with 50 μg/ml fibronectin and 5 μg/ml Bovine Serum (BSA) Alexa Fluor™ 647 conjugate followed by 30 min blocking with 3% BSA, then the non-micropatterned areas where coated with 10 μg/ml GFOGER (Auspep) conjugated to streptavidin using the FastLink Streptavidin Labeling Kit (KA1556, Abnova) according to manufacturer’s instruction.

### Constructing complete atomistic and coarse-grained models

We constructed simulation models that match the protein complexes studied in experiments as accurately as possible. To this end, we built both an atomistic and a coarse-grained Martini 3^44^ model of the bound integrin construct as described here and in the next paragraph. For the fibronectin and antibody-bound integrin structure, we used the PDB id 7NWL, with the bound antibody removed. In this structure the transmembrane and intracellular domains of the integrin alpha and beta are not included. As the starting structure for EGFP, we used the PDB id 2Y0G, where we did not include the three chromatic residues (TYG) in 2Y0G as they are non-standard. To speed up the construction of the complete protein complex, these atomistic structures were individually coarse-grained to the Martini 3 representation with elastic networks and then put together in a single box (^45^; https://github.com/marrink-lab/vermouth-martinize). The box was solvated with default Martini 3 water at 150 mM NaCl. In this coarsegrained representation, the EGFP C-terminus was pulled to the integrin alpha N-terminus (constant rate at 1 nm ns^-1^ with a harmonic force constant of 1000 kJ mol^-1^ nm^-2^) to reach a structure where the termini were less than 1 nm apart. When this state was achieved, the coarsegrained structure (the fibronectin-integrin-EGFP complex) was backmapped to an atomistic representation by aligning the atomistic structures to the pulled coarse-grained system using the C-alpha backbone beads for reference. Finally, as the last few residues at the C-terminus of the EGFP (2Y0G) structure were missing (LGMDELYK), they were added together with the linker using the Modeller loop protocol^46^, i.e., the extended EGFP C-terminus was connected to the N-terminus of the integrin alpha subunit of 7NWL. This final atomistic model contained all sugars and bound ions from the crystal structures and was used as the starting structure in atomistic molecular dynamics simulations. Further, this final atomistic structure of the complete complex was coarse-grained to the Martini 3 representation, which was used as the starting structure in the construction of the complete coarse-grained simulation model (see the next paragraph).

### Constructing the production coarse-grained model

For the Martini 3 coarse-grained production simulations the sugars and manganese cations in the 7NWL structure were not taken into consideration. To maintain the folded structure, an elastic network was added using Martinize2 based on the ElNeDyn protocol^45^. Both intra- and intermolecular contacts were stabilized by the elastic network, but no harmonic bonds were added between EGFP and integrins/fibronectin. The elastic bonds in the linker residues were removed completely. The complex was solvated using insane in default Martini 3 water with 150 mM NaCl^47^ with a minimum periodic distance of 4 nm.

### Simulations of the production coarse-grained models

To run the coarse-grained simulations, we used the GROMACS 2021.2 package^48^. For energy minimization the steepest descent algorithm was used, and during equilibration the default Martini settings were employed, making use of a 1 fs time step up to the point that numerical stability was achieved^49^ The Verlet cut-off scheme was used with a 1.1 nm cut-off for both the Coulombic (reaction-field) and van der Waals interactions. We used v-rescale for the thermostat at 300 K, coupling the protein and solvent in separate groups. Pressure coupling was initially performed using the Berendsen barostat^50^ for isotropic systems at 1 atm. The production runs made use of a 20 fs time step, where the pressure coupling was switched to Parrinello-Rahman^51^. For the pulling simulations, the pull code as implemented in GROMACS 2021.2 was used. The umbrella pulling method was employed to pull EGFP along a vector joining the center of mass of EGFP towards the fibronectin binding site. For each pulling simulation, a rate of 0.1 nm ns^-1^ was used with a harmonic force constant of 1000 kJ mol^-1^ nm^-2^. The videos were made with the VMD movie maker plugin^52^. The production runs spanned 80 ns for the binding site pulling and 3200 ns for the subsequent (non-biased) relaxation.

### Simulations of the atomistic models

To run the atomistic simulations, we used the GROMACS 2021.2 package^48^. The protein was solvated in the presence of 150 mM of sodium chloride at 310 K temperature with a pressure of 1 atmosphere. The LINCS algorithm^53^ was used to constrain the bond lengths in the system during simulations. The CHARMM36m force field^54^ was used to derive the parameters for the protein and the ions. The CHARMM water model^55^ was used to obtain parameters for the water molecules used to solvate the protein. The particle mesh Ewald technique^56^ was used to calculate electrostatic interactions within the simulation system with a real-space cut-off of 1.2 nm. The protein structure was first energy minimized and then subjected to a 100 ns equilibration. Ten independent simulations were then performed for generating the production runs. A time step of 4 fs was used for the simulations using the hydrogen mass partitioning method^57^ For the pulling simulations, the pull code as implemented in GROMACS 2021.2 was used. The umbrella pulling method was employed to pull the EGFP along a vector joining the center of mass (COM) of EGFP towards the fibronectin binding site. For each pulling simulation, a rate of 0.1 nm/ns was used. The videos were made with the VMD movie maker plugin^52^.

### CLEM analyses

U2OS cells were transfected with RUSH-α5 plasmid and cultured from this point grown in medium containing 2.5 mg/ml streptavidin. After 24 hours the cells were split and seeded on fibronectin-coated, photo-etched coverslips (Electron Microscopy Sciences, Hatfield, USA). The cells were left to adhere for 4 hours and incubated for two hours with 250 nM MitoTracker Red CMXRos (M7512, ThermoFisher Scientific). RUSH-α5 was released from the ER upon incubation with medium supplemented with 3 mM biotin for 10 minutes, before fixation in 4% formaldehyde/0.01% glutaraldehyde/PHEM buffer (60 mM PIPES, 25 mM HEPES, 2 mM MgCl2, 10 mM EGTA, pH 6.9), for 1 hour. The coverslips were washed with PHEM buffer, mounted with Mowiol containing 1 μg/ml Hoechst 33342 and examined with a Zeiss LSM780 confocal microscope (Carl Zeiss MicroImaging GmbH, Jena, Germany). A Z-stack covering the whole cell volume of cells expressing distinct RUSH-α5-positive vesicles was acquired using the 63x objective. The relative positioning of the cells on the photoetched coverslips was determined by taking a low magnification DIC image using the 20x objective. After imaging the coverslips were removed from the object glass, washed with PHEM buffer, and fixed in 2% glutaraldehyde/PHEM for 1 hour. Postfixation was done in 1% OsO_4_ and 1.5% KFeCN in the same buffer. Samples were further stained en bloc with 4% aqueous uranyl acetate for 1 h, dehydrated in graded ethanol series and embedded with Epon-filled BEEM capsules (EMS; Polysciences, Inc., 00224) and placed on top of the Mattek dish. After polymerization blocks were trimmed down to the regions previously identified on the confocal microscope and now imprinted on the Epon block. Serial sections (600 nm) were cut on an Ultracut UCT ultramicrotome (Leica, Germany) and collected on formvar-coated slot grids. For observations we routinely used the first section on the block representing the region closest the substrate. Samples were observed in a Thermo Scientific™ Talos^TM^ F200C microscope at 200 kV using a bright field detector for STEM (scanning transmission electron microscopy) imaging. The scanning beam was set to 0.59 mrad angle and camera length was 530 mm. For STEM tomography image series were taken at −56° to 56° tilt angles with 2° increment with a pixel size of 3.18 nm. Tomograms were computed using weighted back projection using the IMOD package. Display of tomogram slices was also performed using IMOD software version 4.9.3. Image overlay of immunofluorescence images and electron micrographs was performed manually using Adobe Photoshop in overlay mode with mitochondria as landmarks.

### Computational clutch model

The clutch model used considers how force transmitted from myosin motors to the substrate is applied to talin molecules and integrin-substrate bonds. Integrins bind and unbind from the substrate through binding rate *k_on_* and unbinding rate *k_off_*, and talin folds and unfolds with folding and unfolding rates *k_fold_* and *k_unfold_. K_off_, k_fold_*, and *k_unfold_* depend on force as previously described experimentally. Binding sites on the substrate are modelled explicitly, whereas integrins are modelled implicitly via a given integrin density *d_int_*. Each time that talin unfolds an adhesion reinforcement event is assumed to happen, which is modelled as an increase in integrin density *d_add_*. Model code and all parameters were taken from previous work^7^. The only differences were the following:

- Our previous work considered that integrin density could both increase (when talin unfolds) and decrease (when integrins unbind from the substrate without talin unfolding). Here, we are only modelling adhesion growth, so we only consider growth.
- We set an upper limit to integrin density (3 times the initial value), to consider that adhesions only grow to a maximum size.
- We decreased the parameter *d_add_* to match the timescale of adhesion growth (to 0.01 or 0.005 integrins/μm^2^).

### Statistical analysis

The GraphPad Prism software was used and the names and/or numbers of individual statistical tests, samples and data points are indicated in figure legends. Unless otherwise noted, all results are representative of three independent experiments and P values <0.05 are shown in graphs.

## Acknowledgements

We thank P. Laasola, J. Siivonen and M. Miihkinen for technical assistance and scientific discussion and the Ivaska lab for critical reading and feedback on the manuscript. B. Goud is acknowledged for hosting M.L.’s research visit at Institut Curie. The Cell Imaging and Cytometry Core (Turku Bioscience Centre, University of Turku) and the University of Helsinki Genome Biology Unit, both supported by Biocenter Finland, and the Euro-BioImaging Finnish Node (Turku Finland) are acknowledged for services, instrumentation, and expertise. We also acknowledge CSC – IT Center for Science for providing computing resources.

This study has been supported by the Academy of Finland (325464 J.I., 331349, 335527, 346135 I.V.), the Academy of Finland CoE in Cell and Tissue Mechanics (J.I.), the Sigrid Juselius Foundation (J.I., I.V.), the Finnish Cultural Foundation (J.L. and E.P), the Finnish Cancer Organization (J.I.), Turku Doctoral Programme of Molecular Medicine (TuDMM; M.L.), Svenska Kulturfonden (M.L.), Victoriastiftelsen, K. Albin Johanssons Stiftelse (M.L.), (Marie Curie Skłodowska fellowship scheme, 101033606 S.K.), Helsinki Institute of Life Science (HiLIFE) Fellow Program (I.V.), Human Frontier Science Program (RGP0059/2019, I.V.), Norwegian Cancer Society (NMP), Spanish Ministry of Science and Innovation (PID2019-110298GB-I00 P. R.-C.), the European Commission (H2020-FETPROACT-01-2016-731957 P.R-C.), The prize “ICREA Academia” for excellence in research P.R.-C., Fundació la Marató de TV3 (201936-30-31 to P.R.-C.), and “la Caixa” Foundation (grant LCF/PR/HR20/52400004 P.R.-C). IBEC is a recipient of a Severo Ochoa Award of Excellence from MINCIN. This study was supported by the Academy of Finland (338537 to GJ), the Sigrid Juselius Foundation (GJ), the Cancer Society of Finland (GJ), the Åbo Akademi University Research Foundation (GJ, CoE CellMech) and the Drug Discovery and Diagnostics strategic funding to Åbo Akademi University (GJ)

## Author contributions

Conceptualization: J.I., M.L, S.M. Methodology: A.B., G.B., J.I., M.L, S.M., F.P., P.R-C., I.V. Formal Analysis: A.B., B.M.H.B., M.C., J.I., G.J., S.K., M.L, M.M. Investigation, B.M.H.B., M.C., J.I., S.K., M.L. N.M.P, L.M, M.M., P.R-C. Resources: A.B., G.B., G.J., F.P., I.V. Visualization: A.B., H.H., M.L, M.M., Writing – Original Draft: J.I., M.L. Writing - review & editing: H.H., J.I., M.L., M.M., Supervision: A.B., J.I., S.M., I.V. Funding Acquisition: J.I., M.L., M.M.

---

**Extended Data Figure 1.**
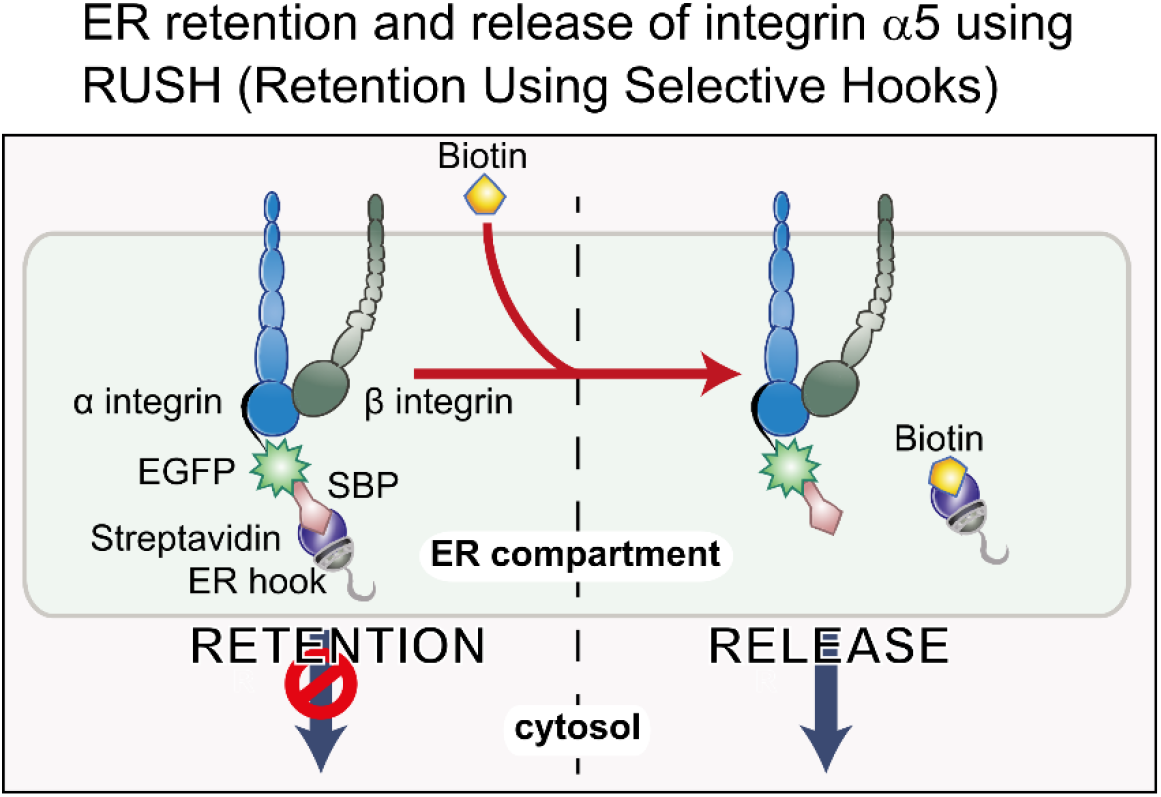
Principle of the RUSH system applied to integrin α5. In all experiments SBP-EGFP-ITGA5 (RUSH-α5) is co-expressed with streptavidin-KDEL (ER hook). In the absence of biotin this combined complex is retained within the ER. Biotin addition displaces the ER hook and releases RUSH-α5 into the cytoplasm.

**Extended Data Figure 2.**
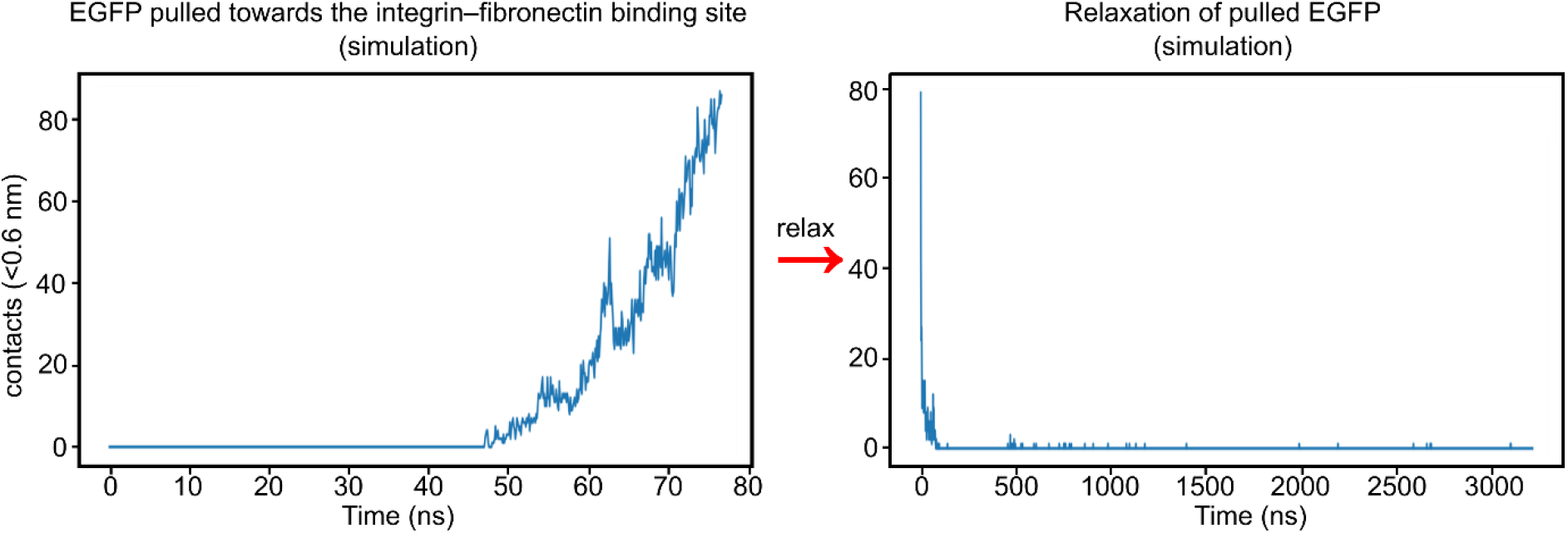
Number of contacts between EGFP and fibronectin during simulations of the coarse-grained model. Left: simulation of EGFP being pulled towards the fibronectin binding site, starting when the C-terminus of the EGFP and the N-terminus of the integrin α5 are less than 1 nm apart, the linker included, leading to the formation of contacts (Movie 2). Right: simulation of a fully stretched EGFP, initially in close proximity to the fibronectin binding site, that is allowed to relax without a biasing force resulting in a spontaneous and rapid loss of contacts (<100 ns; Movie 3). The pulling process spanned 8 nm and 80 ns. The relaxation spanned 3200 ns. Contacts were calculated between EGFP and fibronectin with a cutoff of 0.6 nm.

**Extended data figure 3.**
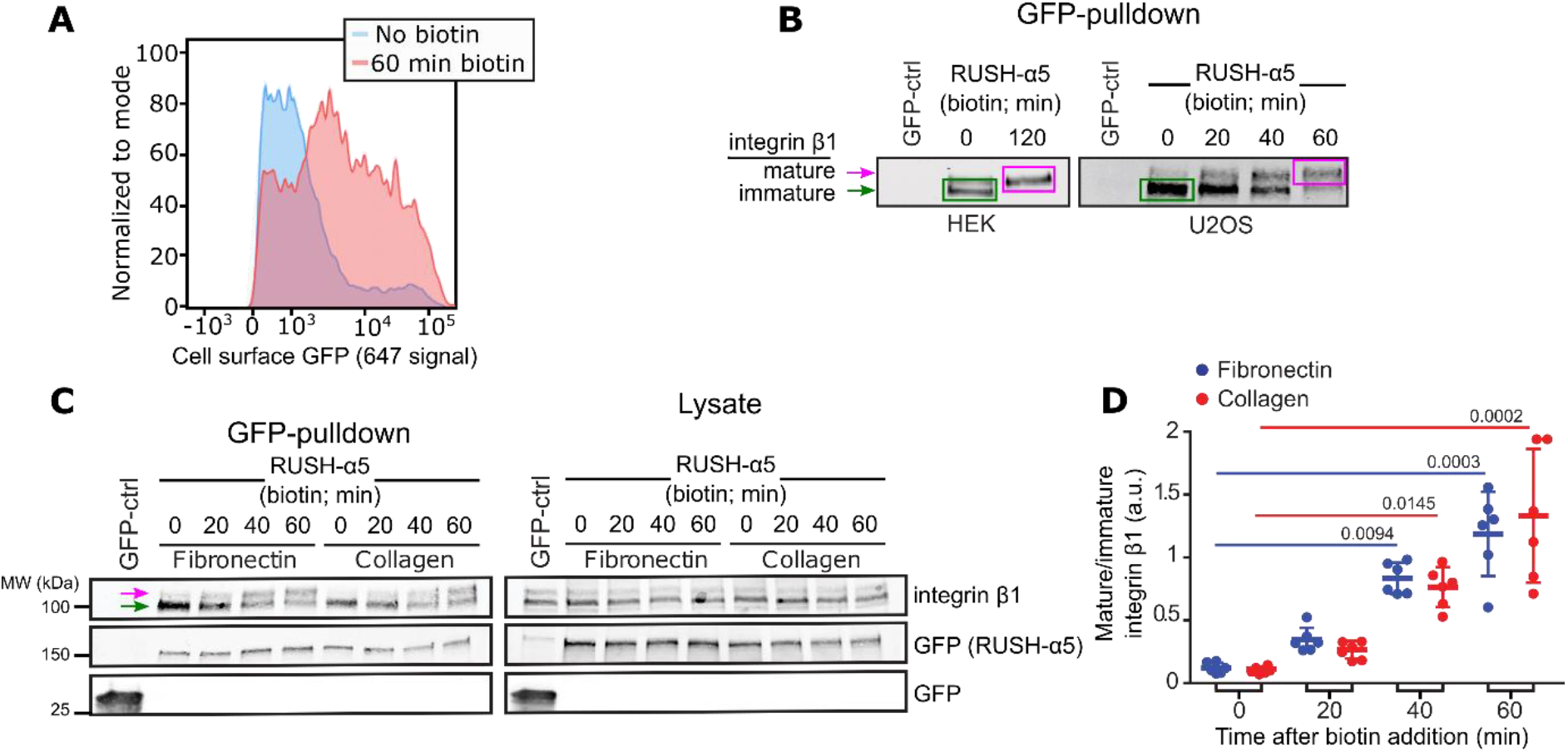
RUSH-α5 is expressed on the cell surface and forms a functional heterodimer with β1-integrin. **a)** Representative flow cytometry analysis of cell-surface RUSH-α5 levels (detected with the anti-GFP-AF647 antibody) in RUSH-α5-expressing U2OS cells ± biotin. **b)** Representative immunoblots of GFP-pulldowns performed in RUSH-α5 or control transfected cells ± biotin treatment for the indicated times and probed for endogenous integrin β1. The faster migrating band of immature integrin β1 is indicated by a green arrow and box and the slower migrating band of mature integrin β1 with a magenta arrow and box. **c)** Representative immunoblot of GFP-pulldowns performed in RUSH-α5 or control transfected cells plated on fibronectin or collagen and probed for endogenous integrin β1 and for GFP. Mature (magenta arrow) and immature (green arrow) integrin β1 are indicated. **d)** Quantification of the relative fraction of mature to immature integrin β1 interacting with RUSH-α5 ± biotin treatment for the indicated times. N= 6 independent experiments; One-way ANOVA, Dunn’s multiple comparison test.

**Extended data Figure 4.**
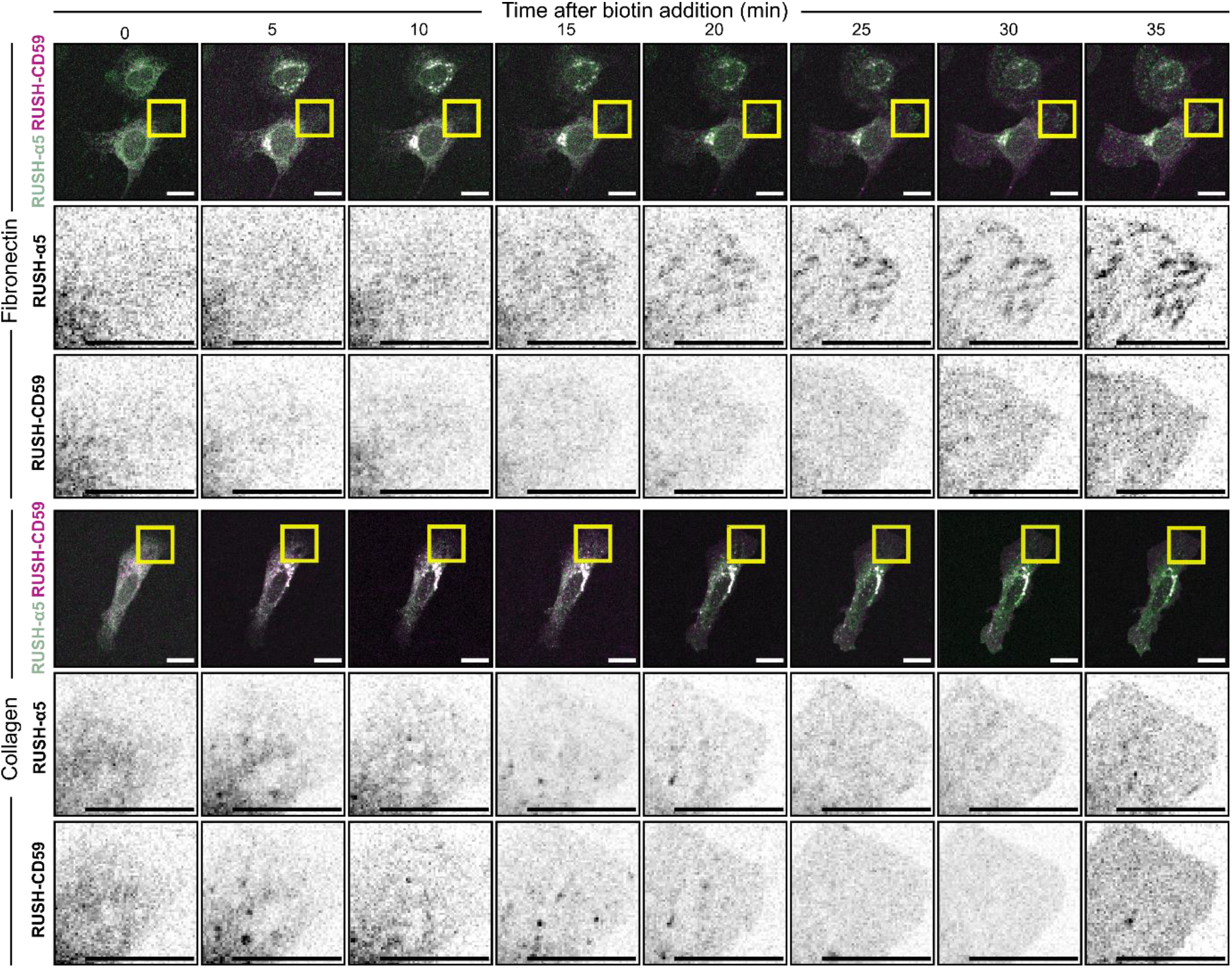
RUSH-α5 recruitment to adhesions is ligand-dependent. Representative images (see Movie 8) of U2OS cells co-expressing RUSH-α5 and RUSH-CD59 and plated on fibronectin (top) or collagen (bottom) ± biotin treatment for the indicated times. Insets represent ROIs that are magnified. Scale bars: 20 μm.

**Extended data Figure 5.**
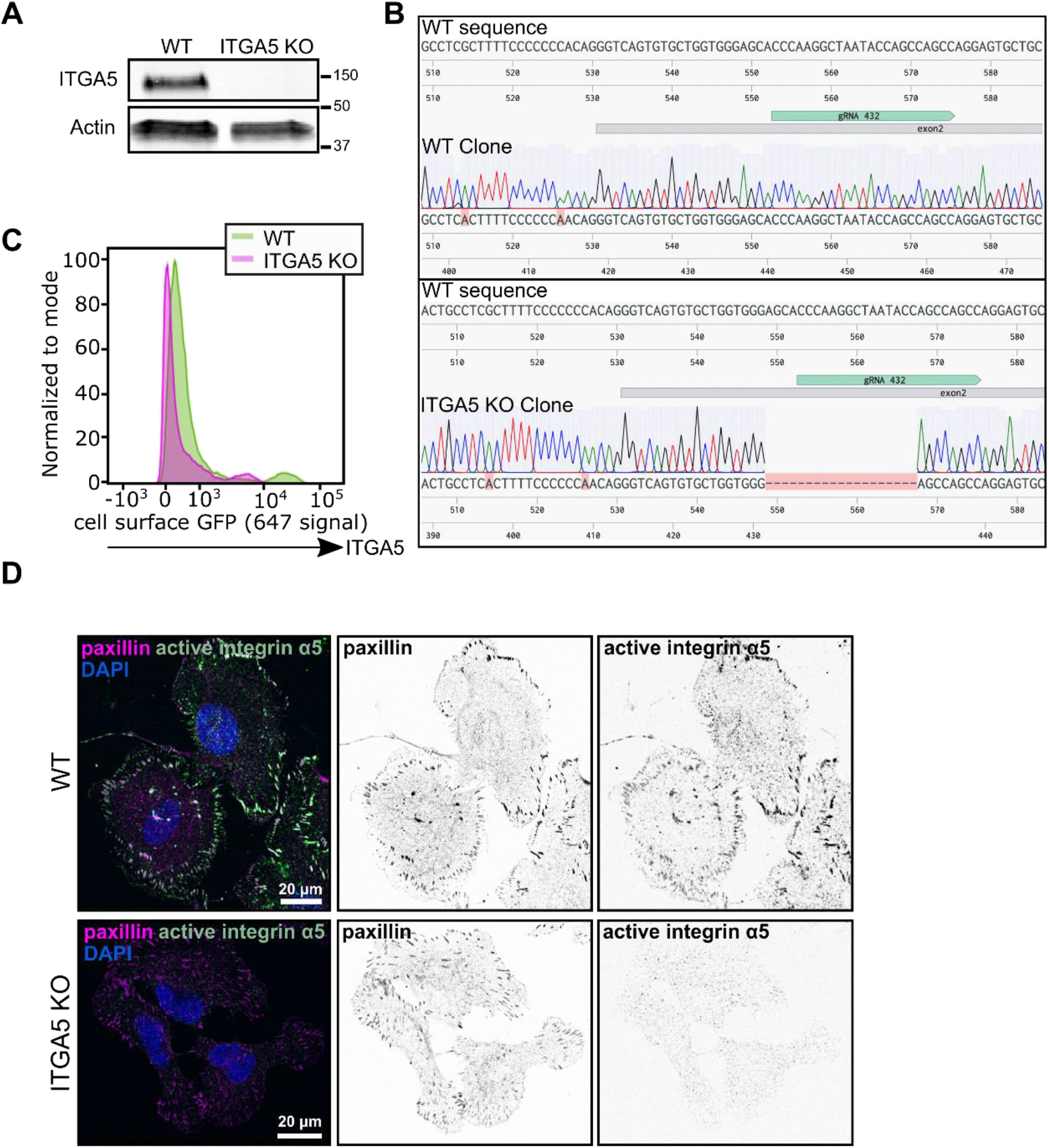
Validation of ITGA5 CRISPR-Cas9 KO U2OS cells. **a)** Western blot analysis of WT and ITGA5 KO single cell clones showing the efficiency of the CRISPR-Cas9 ITGA5 KO in U2OS cells. **b)** Genome sequence alignment of U2OS WT and ITGA5 KO clones with ITGA5 WT sequence. The targeted exon and the gRNA used for CRISPR KO positions are indicated. **c)** Representative flow cytometry analysis of cell surface integrin-α5 in U2OS WT and ITGA5 KO clones. **d)** Images of WT and ITGA5 KO U2OS clones stained for active integrin α5 (SNAKA51) and paxillin. Scale bar: 20μm.

**Extended data Figure 6.**
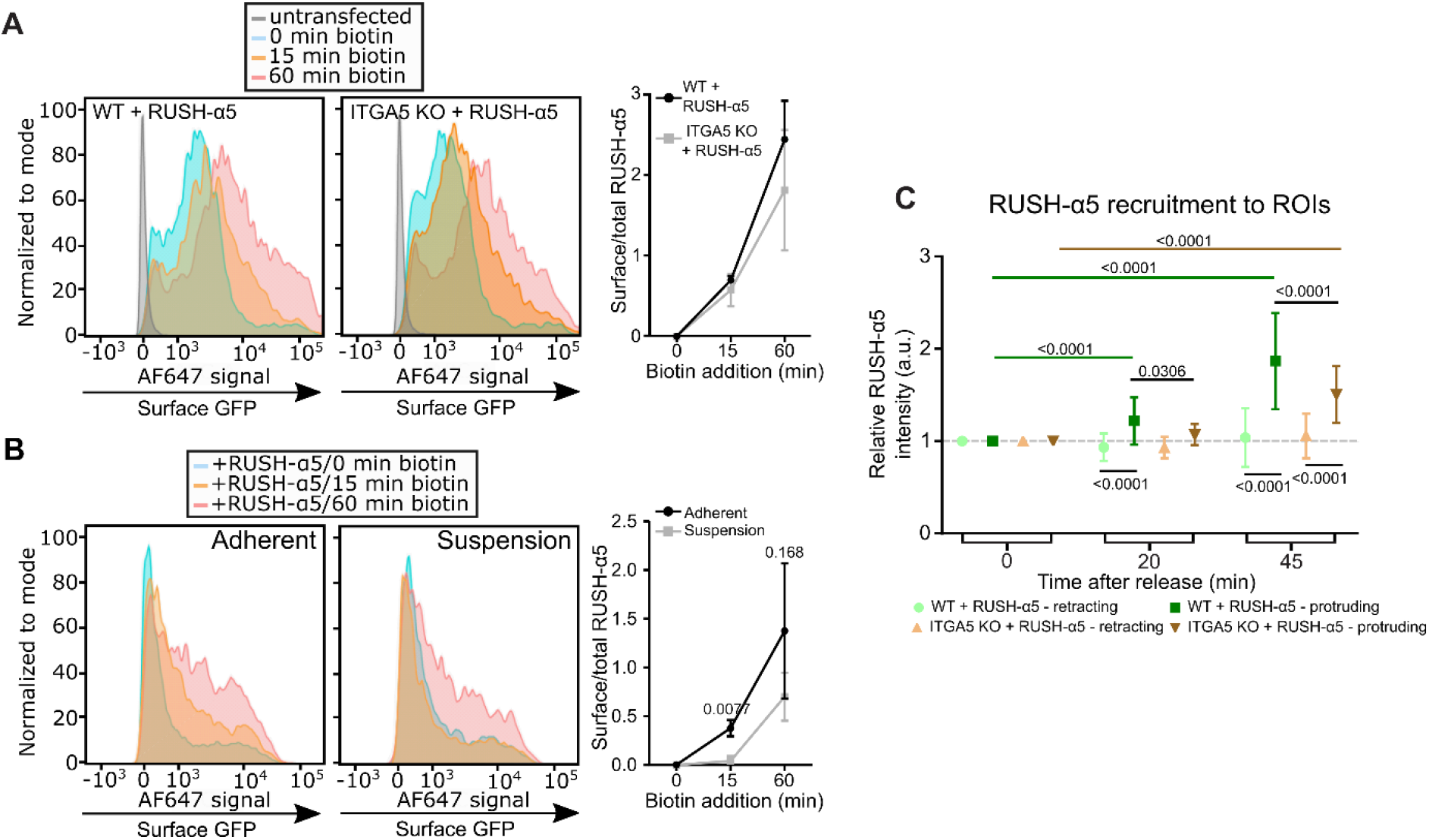
Early release of RUSH-α5 is adhesion dependent and polarized recruitment to protrusion is supported by endogenous integrin α5. **a)** Flow cytometry analysis of cell-surface RUSH-α5 levels (detected with the anti-GFP-AF647 antibody) in WT and ITGA5 KO U2OS cells expressing RUSH-α5 ± biotin treatment for the indicated times. Representative histograms and quantification of cell surface GFP (ratio of the geometric means of the surface signal divided by the total GFP signal, normalized by subtracting the 0 min value) are shown. N=3 independent experiments. **b)** Flow cytometry analysis of cell-surface RUSH-α5 levels in adherent versus suspension cells expressing RUSH-α5 ± biotin treatment for the indicated times. Representative histograms and quantification of cell surface GFP (analysed as in a) are shown. N=3 independent experiments. Two-tailed paired t-test. **c)** Quantifications of RUSH-α5 intensity in ROIs (retracting or protruding areas) in WT and ITGA5 KO U2OS cells ± biotin treatment for the indicated times. One-way ANOVA, Holm-Sidak’s multiple comparison test. Data are mean ± SD; N= 59 WT cells, 53 ITGA5 KO cells, pooled from 3 independent experiments.

**Extended Data Figure 7.**
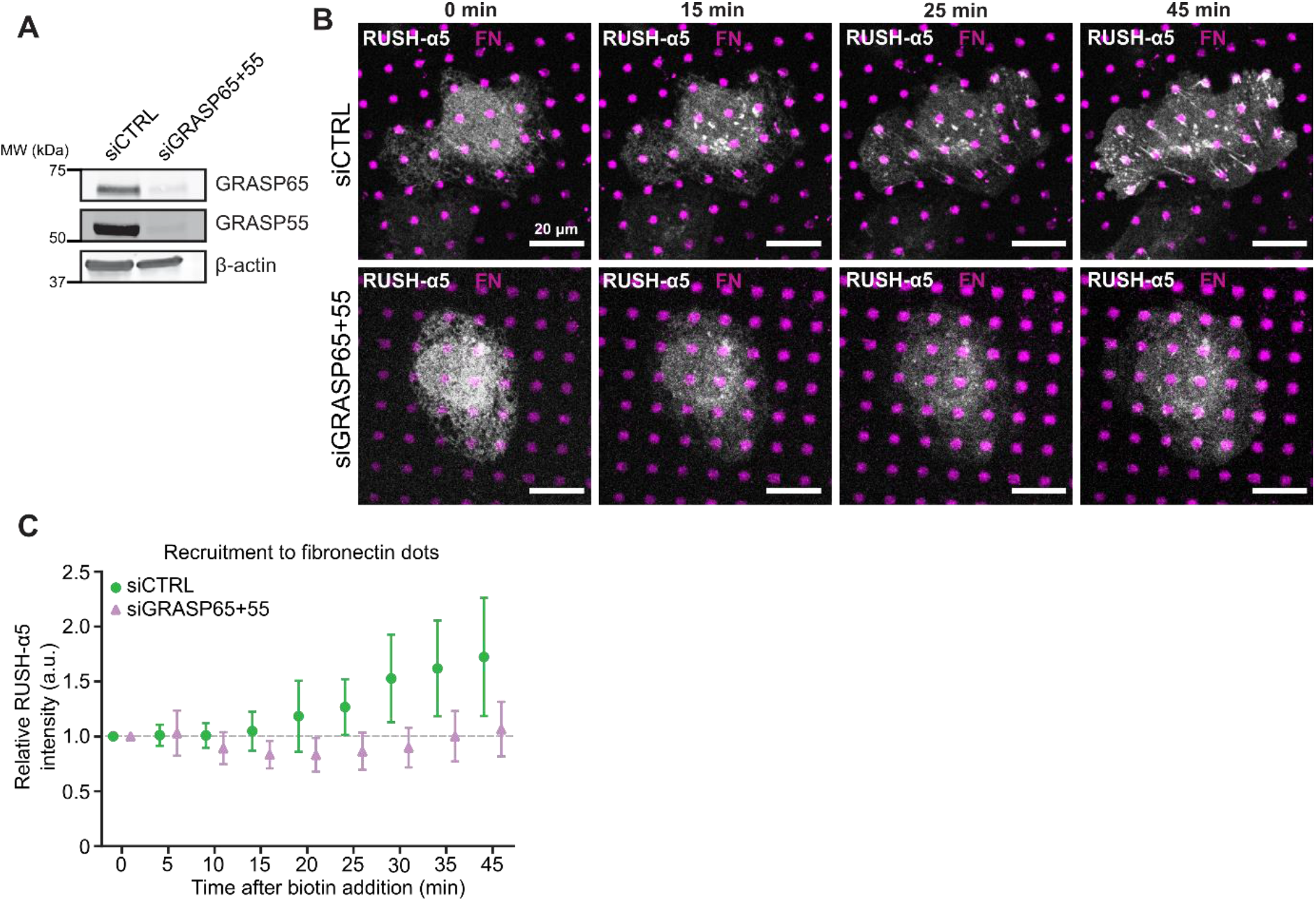
Early release of RUSH-α5 is sensitive to GRASP silencing. **a)** Immunoblot of lysates collected from control-silenced or GRASP65 and GRASP55-silenced cells used in b, c, probed for GRASP65 and GRASP55. β-actin was probed as a loading control. **b)** Representative immunofluorescence images of control-silenced or GRASP65 and GRASP55-silenced cells expressing RUSH-α5 and plated on dual-coated micropatterns (cyan dots, fibronectin; non-fluorescent regions, collagen peptide GFOGER). **c)** Relative recruitment of RUSH-α5 in control-silenced or GRASP65 and GRASP55-silenced cells to fibronectin dots. Data are mean ± SD; n = 9 ctrl cells, 11 siGRASP cells (36 and 44 dots respectively) from one experiment. Scale bars: 20 μm.

**Extended Data Figure 8.**
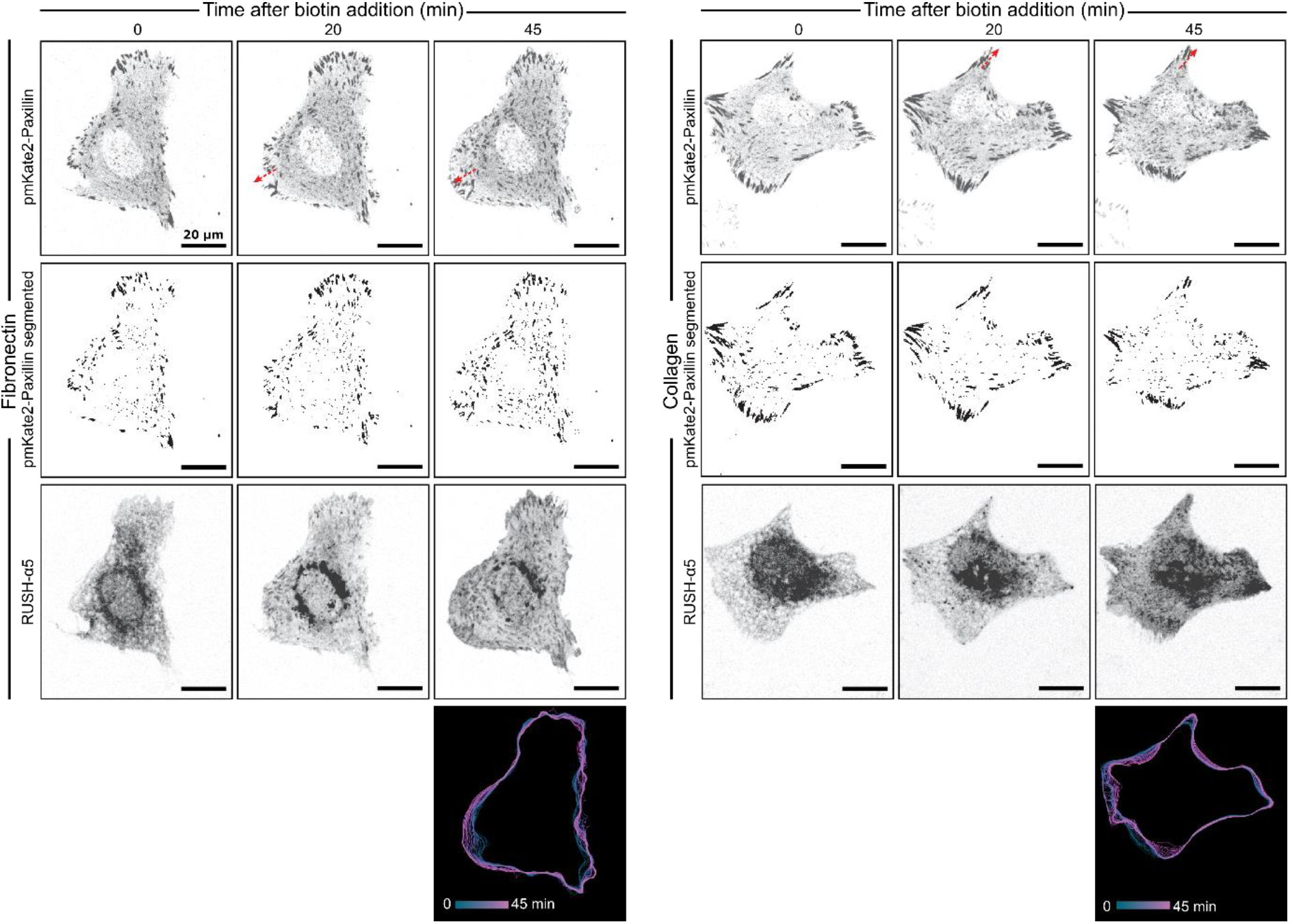
Released RUSH-α5 increases protrusion length on fibronectin. Representative images and trackmaps related to figure 5i. Red arrows indicated direction of adhesion growth.

**Extended Data Table 1.**
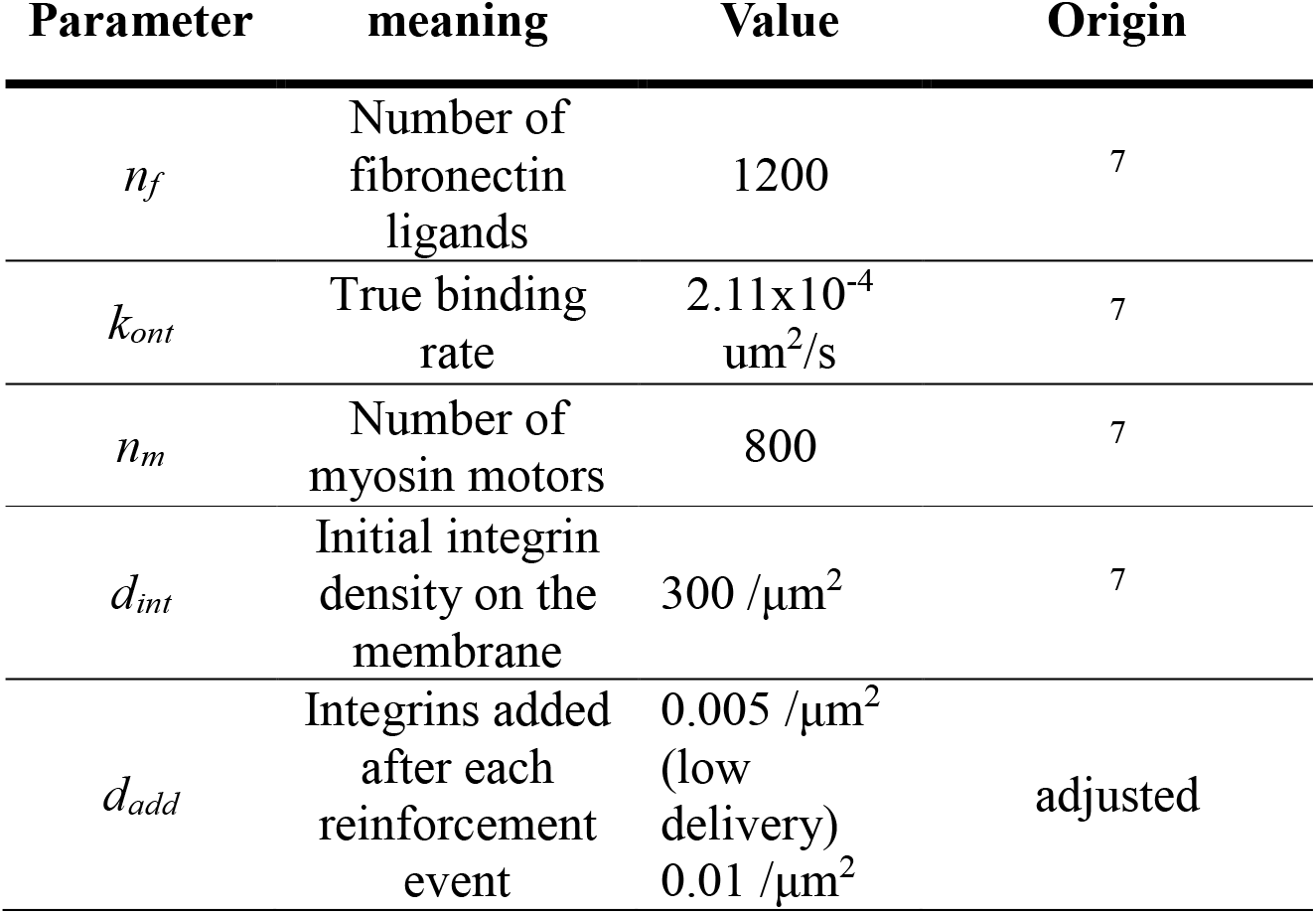
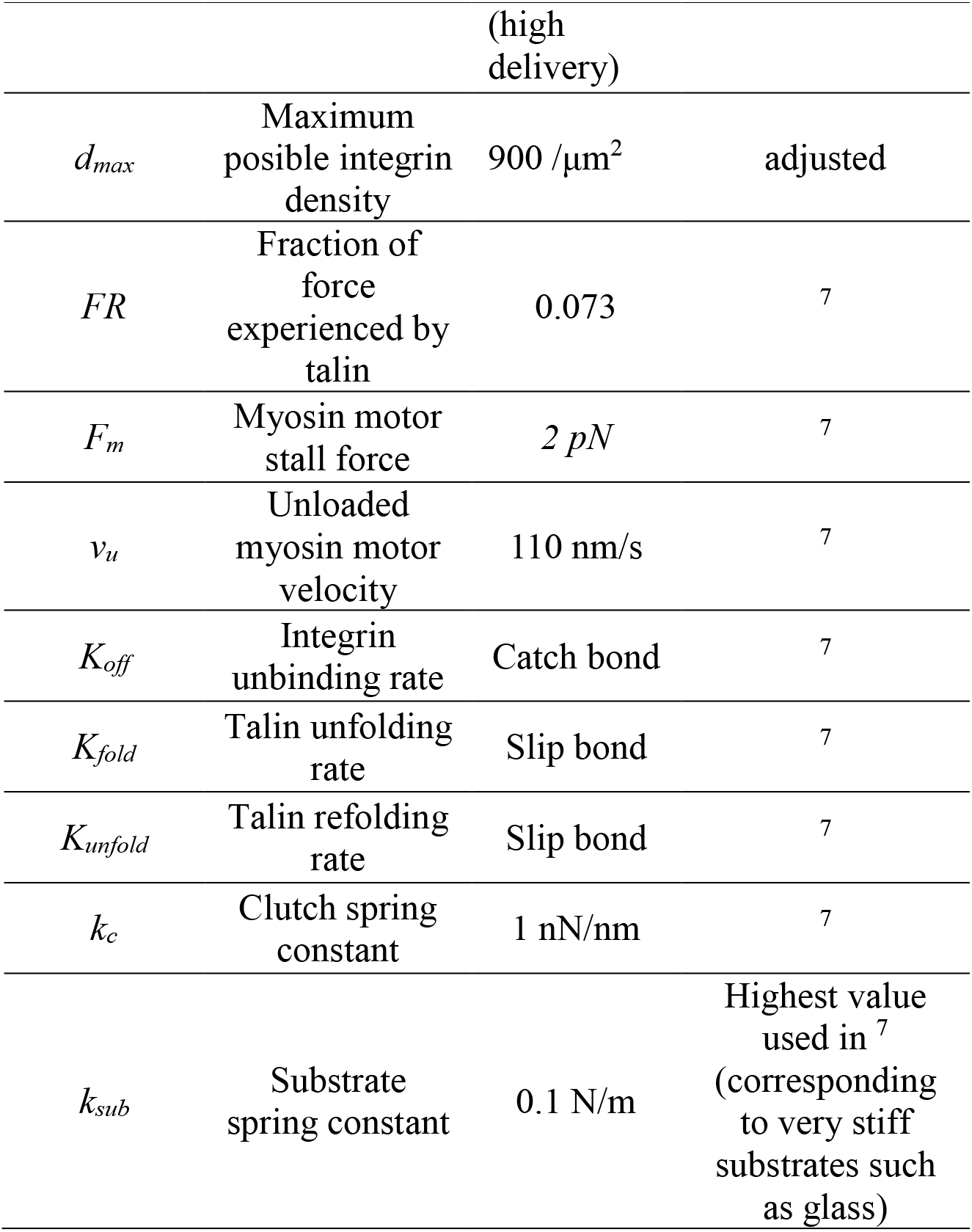
Model parameters. Note that all parameters are taken from except for *d_add_* and *d_max_*.

## Supplementary Movies

**Movie 1.** Turning model of RUSH-α5 (EGFP-integrin α5) - integrin-β1 heterodimer bound to fibronectin based on (PDB: 7NWL) structure of the heterodimer.

**Movie 2**

For the coarse-grained model, the pulling of EGFP towards the fibronectin binding site. The pulling between the EGFP and the fibronectin binding site starts from the situation where the C-terminus of the EGFP and the N-terminus of the Integrin alpha are less than 1 nm apart, including the linker. The movie is divided into three parts. It starts with a full rotation around the initial configuration. Then, the pulling is performed (at a constant rate with 1000 kJ mol^-1^ nm^-2^ at 0.1 nm/ns). Finally, shown is a full rotation around the final state, later used as the first frame for relaxation in Movie 3. The movie repeats itself in reverse. EGFP is blue, the linker is red, and integrin alpha is orange (one molecule). Integrin beta is yellow, and fibronectin is moss green. The pulling process spans 8 nm and 80 ns.

**Movie 3**

For the coarse-grained simulation model, the spontaneous (non-biased) relaxation of the final pulled state of Movie 2. Contacts (< 0.6 nm distance) between the EGFP and fibronectin are indicated with magenta (Extended data Figure 2). EGFP is blue, the linker is red, and integrin alpha is orange (one molecule). Integrin beta is yellow, and fibronectin is moss green. The relaxation spans 3200 ns.

**Movie 4**

Fully atomistic molecular dynamics simulation of the EGFP attached to the α-subunit of the integrin molecule. The movie depicts a demonstrative simulation of the EGFP (in green) bound to the α-subunit (blue) of integrin. The β-subunit is shown in red color. The fibronectin bound to integrin is shown in gray. No additional forces were applied in this simulation. The simulation is 100 ns long.

**Movie 5**

Fully atomistic steered molecular dynamics simulation of the EGFP attached to the α-subunit of the integrin molecule. The video depicts a demonstrative simulation of the EGFP (in green) bound to the α-subunit (blue) of integrin. The EGFP is pulled towards the fibronectin (gray) binding site with a force of 25 kJ/mol/nm^-2^. The β-subunit on integrin is shown in red. The simulation is 100 ns long.

**Movie 6**

Fully atomistic steered molecular dynamics simulation of the EGFP attached to the α-subunit of the integrin molecule. The movie depicts a demonstrative simulation of the EGFP (in green) bound to the α-subunit (blue) of integrin. The EGFP is pulled towards the fibronectin (gray) binding site with a force of 50 kJ/mol/nm^-2^. The β-subunit on integrin is shown in red. The simulation is 100 ns long.

**Movie 7.**

Time lapse imaging of RUSH-α5 (SBP-EGFP-integrin α5)-expressing U2OS cell plated on fibronectin (10 μg/ml), biotin added after acquisition of timepoint 0 min.

**Movie 8.**

Time lapse imaging of U2OS cells co-expressing RUSH-α5 (green) and RUSH-CD59 (magenta) and plated on fibronectin (left, 10 μg/ml) or collagen (right, 10 μg/ml), biotin added after acquisition of timepoint 0 min.

**Movie 9.**

Time lapse imaging of U2OS cells co-expressing RUSH-α5 (green) and the ER marker ERoxBFP (magenta) plated on fibronectin (10 μg/ml), biotin added after acquisition of timepoint 0 min.

